# A Chemoenzymatic Strategy for Site-Specific Glyco-Tagging of Native Proteins for the Development of Biologicals

**DOI:** 10.1101/2024.08.13.607754

**Authors:** Ana Gimeno, Anna M. Ehlers, Sandra Delgado, Jan-Willem H. Langenbach, Leendert J. van den Bos, John A.W. Kruijtzer, Bruno G.A. Guigas, Geert-Jan Boons

**Affiliations:** Chemical Biology and Drug Discovery, Utrecht Institute for Pharmaceutical Sciences, and Bijvoet Center for Biomolecular Research, Utrecht University, 3584 CG Utrecht, The Netherlands; CIC bioGUNE, Basque Research & Technology Alliance (BRTA), Bizkaia Technology Park, Building 800, 48160 Derio, Bizkaia, Spain; EnzyTag BV, Daelderweg 9, NL-6361 HK Nuth, The Netherlands; Leiden University Center of Infectious Diseases, Leiden University Medical Center, Leiden 2333 ZA, The Netherlands; Complex Carbohydrate Research Center, University of Georgia, Athens, Georgia 30602, United States; Department of Chemistry, University of Georgia, Athens, Georgia 30602, United States

**Author notes:** **Corresponding Author:** Geert-Jan Boons Chemical Biology and Drug Discovery, Utrecht Institute for Pharmaceutical Sciences, Utrecht University, Utrecht 3584 CG, The Netherlands; Complex Carbohydrate Research Center, University of Georgia, Athens, Georgia 30602, United States; Bijvoet Center for Biomolecular Research, Utrecht University, Utrecht 3584, The Netherlands; Chemistry Department, University of Georgia, Athens, Georgia 30602, United States.

## Abstract

Glycosylation is an attractive approach to enhance biological properties of pharmaceutical proteins, however, precise installation of glycans for structure-function studies remains challenging. Here, we describe a chemoenzymatic methodology for glyco-tagging of proteins by peptidoligase catalyzed modification of the *N*-terminus of a protein with a synthetic glycopeptide ester having an *N*-acetyl-glucosamine (GlcNAc) moiety to generate a *N*-GlcNAc modified protein. The GlcNAc moiety can be elaborated into complex glycans by *trans*-glycosylation using a well-defined sugar oxazolines and mutant forms of endo β-*N*-acetylglucosaminidases (ENGases). The glyco-tagging methodology makes it possible to modify *on-demand* therapeutic proteins, including proteins heterologously expressed in *E. coli,* with diverse glycan structures. As a proof of principle, the *N*-terminus of interleukin (IL)-18 and interferon (IFN)α-2a was modified by a glycopeptide harboring a complex *N*-glycan without compromising biological potencies. The glyco-tagging methodology was also used to prepare several glycosylated insulin variants that exhibit reduced oligomerization, aggregation and fibrillization yet maintained cell signaling properties, which is attractive for the development of insulins with improved shelf-lives. It was found that by employing different peptidoligases, it is possible to modify either the A or both chains of human insulin.

## INTRODUCTION

Biologicals are a fast-growing class of pharmaceuticals that accounted for approximately a third of drugs approved by the FDA in 2022.^1,2^ However, native peptides and proteins often exhibit poor pharmacokinetic profiles (PK) undergoing rapid proteolytic degradation or clearance. PEGylation is a widely applied strategy to improve PK profiles and biological storability of biologics and to date thirty-eight PEGylated therapeutics have been approved by the FDA, including growth factors, Erythropoietin (EPO), interferons and coagulation factors.^3^ *N*-terminal site-specific PEGylation is attractive an provides therapeutics with improved PK profiles with minimal interference of the protein’s secondary structure and biological activity.^4^ Safety concerns such as PEG hypersensitivity, immunogenicity and bioaccumulation have, however, been noted^5,6^ and other pegylated products have failed clinical testing such as PEGylated insulin (Lispro) due to hepatic toxicity.^7^ As an alternative strategy, glycosylation, which is a common post-translational modification,^8^ can be exploited to enhance properties of biopharmaceuticals. In fact, the incorporation of carbohydrate polymers such as dextran or polysialylation has received considerable attention.^9^ Glycosylation impacts protein folding, stability, and proteolytic degradation and can substantially improve circulation times.^10,11^ Glycosylated variants of proteins such as human growth hormones,^12^ insulin^13–18^ and glucagon-like peptide-1^19,20^ exhibit improved therapeutic profiles compared to their non-glycosylated counterparts. Similarly, fusion proteins bearing an additional glycosylated peptide domain have also been described with improved PK profiles.^21,22^ For example, Corifollitropin alfa (Elonva®) is a recombinant follicle-stimulating hormone (FSH) analog that is composed of the β-subunit of FSH fused to the C-terminal peptide of the human chorionic gonadotropin (hCG) β-subunit containing four *O*-linked glycosylation sites. This chimeric protein has a 4-fold increase in elimination half-life and enhanced *in vivo* bioactivity compared to wild-type FSH.^23–25^ Thus, the assembly of fusion proteins tagged with a peptide sequence containing one or more glycosylation sites is an attractive strategy to improve properties of biologicals.

Despite advances, controlled modification of proteins with glycans remains challenging, thereby complicating structure-function studies. Glycan structures are not precisely defined at the genetic level and as a result eukaryotic expression systems generally provide mixtures of different glycoforms.^26^ This hurdle may be overcome by synthetic or semi-synthetic approaches,^27–31^ although current methodologies to modify native proteins with specific glycans often lack selectivity leading to heterogeneity.^32^

Here, we describe a chemoenzymatic methodology that makes it possible to modify the *N*-terminus of native proteins with well-defined glycopeptide tags without introducing non-natural modifications.^33^ Specifically, the *N*-terminus of a protein is ligated with a synthetic glycopeptide ester having an *N*-acetyl-glucosamine (GlcNAc) moiety to generate a *N*-GlcNAc modified protein (Figure 1). The ligation is catalyzed by peptidoligases, which are genetically modified versions of subtilisin from *B. subtilis,* wherein a serine in the active site is exchanged with cysteine, thereby abolishing hydrolytic activity.^34,35^ Their active sites have been engineered to accommodate a wide range of amino acid side chains at the C- and N-terminal ligation junction, resulting in enzymes with broader substrate specificity. Next, the GlcNAc moiety of the resulting glycoprotein can be elaborated into complex glycans by *trans*-glycosylation using mutant forms of endo β-*N*-acetylglucosaminidases (ENGases).^36–38^ These mutant enzymes lack natural hydrolytic activity and instead can use activated glycan oxazolines as donor substrates to install complex glycans onto *N*-GlcNAc moieties. The glyco-tagging methodology described here enables modifying *on-demand* therapeutic proteins, including proteins expressed in *E. coli,* with diverse glycan structures. As a proof of principle, we employed the approach to modify the *N*-terminus of cytokines Interleukin-18 (IL-18) and interferon alpha-2a (IFNα-2a) by a glycopeptide harboring a complex *N*-glycan and demonstrated that the modification does not affect biological potencies. The methodology was also employed to prepare several glycosylated insulin variants that exhibit reduced oligomerization, aggregation and fibrillization yet maintaining cell signaling properties. By employing different peptidoligases, it was possible to modify either the A or both chains of human insulin.

**Figure 1.**
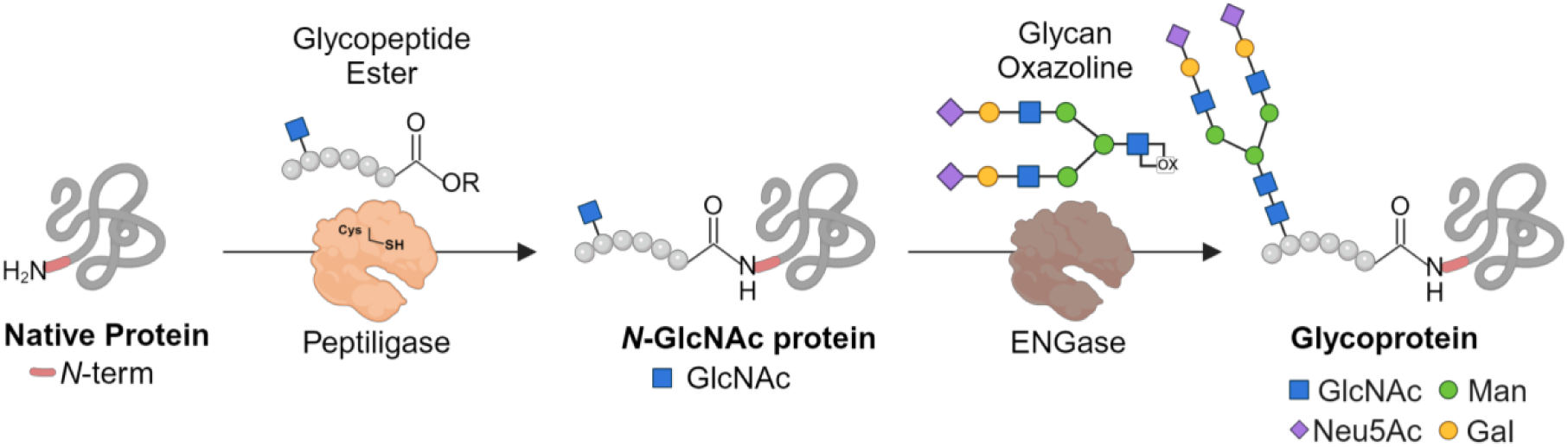
Chemoenzymatic strategy for specific glyco-tagging of native proteins.

## RESULTS AND DISCUSSION

### Chemical Synthesis of a Glycopeptide Ester and Model Ligation

First, a methodology was developed to prepare glycopeptide esters having an *N*-glycosylation site. The resulting compound was employed for a ligation with a model peptide substrate using omniligase-1, which is a readily available and well-characterized peptidoligase.^39,40^ As glycopeptide ester, we selected compound **1** which was ligated with peptide **2** to give glycopeptide **3** (Figure 2A). The P4-P1 residues of the acyl donor are important for ligation efficiency (Figure 2B), and therefore **1** incorporates amino acids at these positions that are preferred by omniligase-1.^41^ To prevent that the glycosylation site of the glycopeptide ester interferes with the activity of the peptidoligase, it was positioned outside the binding pocket of the enzyme. Furthermore, it contains a typical amino acid sequence for *N*-glycosylation (N-X-S/T) to provide a natural *N*-glycan sequon. In particular, the N-A-T sequence was selected because of its high abundance in natural glycoproteins.^42^ To avoid any further reactivity at glycopeptide **1**, the *N*-terminal α-amine was capped by acetylation. The *N*-terminal amino acid sequence of the acyl acceptor is also important for ligation efficiency. Omniligase-1 prefers small apolar amino acids at positions P1’ and P2’, and therefore the *N*-terminus of **2** has an Ala-Ala sequence at these positions (Figure 2B).

**Figure 2.**
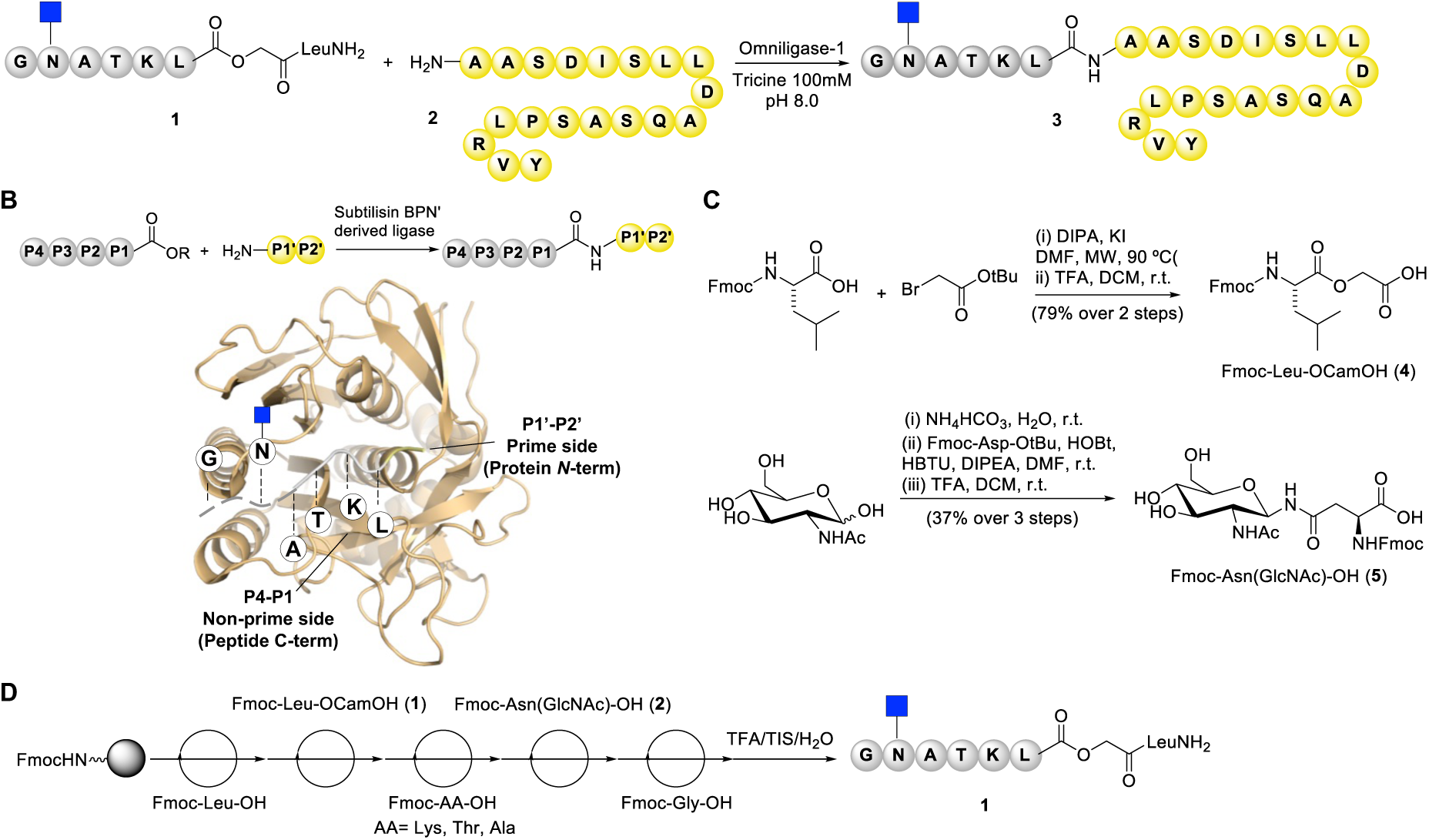
Synthesis and reactivity assays of glycopeptide ester **1**. A) Reaction between glycopeptide **1** and an *N*-terminal di-alanine model peptide **2**. The ligation product **3** was obtained in a 74% isolated yield referred to glycopeptide **1**, used as limiting reagent. Reaction was performed using 2:1 ratio of peptide **2** to glycopeptide **1** in the presence of 1.5 mol% enzyme loading. B) Peptide ligation catalyzed by subtilisin-derived ligases. Below: structure of subtilisin BPN’ (pdb 1SBN) with a schematic representation of P4-P1 and P1’-P2’ positions of the acyl donor and acyl acceptor fragments respectively, which are important for successful peptide ligation. C) Chemical synthesis of building blocks **4** and **5**. D) Solid phase peptide synthesis of glycopeptide ester **1**.

The ester linkage of **1** was installed using Fmoc-Leu-OCH_2_COOH (**4**, Cam Ester) as building block (Figure 2C), which could readily be prepared by condensation of Fmoc-protected leucine with tert-butyl bromoacetate followed by the removal of the *t*-butyl ester using TFA in DCM (79% overall yield). The glycosylated amino acid Fmoc-Asn(GlcNAc)-OH (**5**) was employed to install the carbohydrate moiety of **1**, which was prepared by Kochetkov amination of *N*-acetyl glucosamine followed by coupling of the resulting α-glycosylamine with *N*-α-Fmoc-protected L-aspartic acid-tert-butyl ester and then by treatment with TFA/DCM to remove the *t*-butyl ester (Figure 2C).^43^ The GlcNAc-containing glycopeptide ester **1** was synthesized on a Rink Amide AM LL resin using a CEM Liberty 12-channel automated microwave peptide synthesizer employing a standard Fmoc-solid-phase peptide synthesis protocol (Figure 2D). The glycopeptide ester was cleaved from the resin with simultaneous removal of the side chain protecting groups of the amino acids by treatment with TFA/TIS/water and then purified by HPLC using a C18 reverse column. Mass spectrometry and NMR confirmed the structural integrity of the compound. Ligation of glycopeptide ester **1** with peptide **2** in the presence of omniligase-1 proceeded smoothly in 100 mM tricine at pH 8.0 to provide the corresponding glycopeptide **3** in a 74% yield after purification by C18 reverse phase HPLC (Figure 2A).^44^ A by-product was also isolated in which the ester of **1** had been hydrolyzed.

### Ligation of Glycosylated Peptide Esters with Proteins

Encouraged by the successful synthesis of a glycopeptide **3** by peptide ligation, attention was focused on the modification of proteins with a glycopeptide tag. First, the ligation of glycopeptide ester **1** with the carbohydrate binding domain of human Galectin-3 (*h*Gal3-CRD), which can easily be expressed in *E. coli,*^45^ was investigated. The reaction was conducted at 100-150 mM protein concentration with 1 mol% omniligase-1 and analyzed by mass spectrometry after 2 h (Table S5 for additional details), which indicated a relatively low conversion of ∼20%. *h*Gal3-CRD has the non-polar residues Met-Leu at the *N*-terminus, which is sub-optimal for omniligase-1, at least under the employed reaction conditions. Therefore, we investigated thymoligase as peptide ligase, which was developed to accept negatively charged amino acids at position P1’.^46^ Interestingly, the use of this ligase resulted in an improved conversion of ∼50%, demonstrating its compatibility with other amino acids, including those with amphipathic characteristics such as methionine.

Next, attention was focused on glyco-tagging of the biomedically important cytokines IL-18 and IFNα-2a. Cytokines are small signaling proteins that can modulate processes such as inflammation and immune response. They are emerging as attractive therapeutics for various immune-related diseases, however, their short half-live limit their application and methodologies are needed to increase the stability of these proteins.^47^ IL-18 is experiencing a renascence in cancer therapy by enhancing IFN-γ production by tumor-infiltrating T cells^48^ and IFNα-2a is an established therapeutic for the treatment of Hepatitis B and C (brand name of pegylated form: Pegasys) due to its anti-proliferative capacity on virus-infected immune cells.^49^ Both proteins were recombinantly expressed in *E. coli*^50,51^ as soluble entities (Figures S6 and S7). A TEV (tobacco etch virus) protease cleavage site (ENLYFQ/G) was introduced into the respective synthetic gene constructs to expose amino acids other than methionine, which serves as the initiation translating residue in *E. coli,* at the *N*-terminus of the proteins. After TEV cleavage, the IL-18 and IFNα-2a sequences incorporate *N*-terminal Gly, which small and hydrophobic character would enhance the efficiency of ligation.^41^ In the case of IFNα-2a, which natural sequence starts with a Cys involved in a disulfide bond, an additional apolar residue was included so that the final protein contained a Gly-Phe moiety at the *N*-terminus. The integrity of the expressed proteins was confirmed by intact protein MS after TEV cleavage and purification by affinity chromatography (Figures 3A and 3B). The ligation between IL-18 (**6**) or IFNα-2a (**7**) and glycopeptide **3** (5 eq.) was performed with thymoligase and provided the corresponding GlcNAc variants **6a** and **7a**, respectively (Figure 3C). The employed thymoligase contained a His tag and could therefore be removed by Ni-NTA affinity chromatography. Analysis of the protein fractions by intact protein mass spectrometry showed predominant species corresponding to the ligation of glycopeptide **3** to IL-18 and IFNα-2a with peaks with m/z at 23748.47 Da and 20271.38 Da, respectively (Figures 3D and 3E). A small amount of unmodified protein was detected in each case.

**Figure 3.**
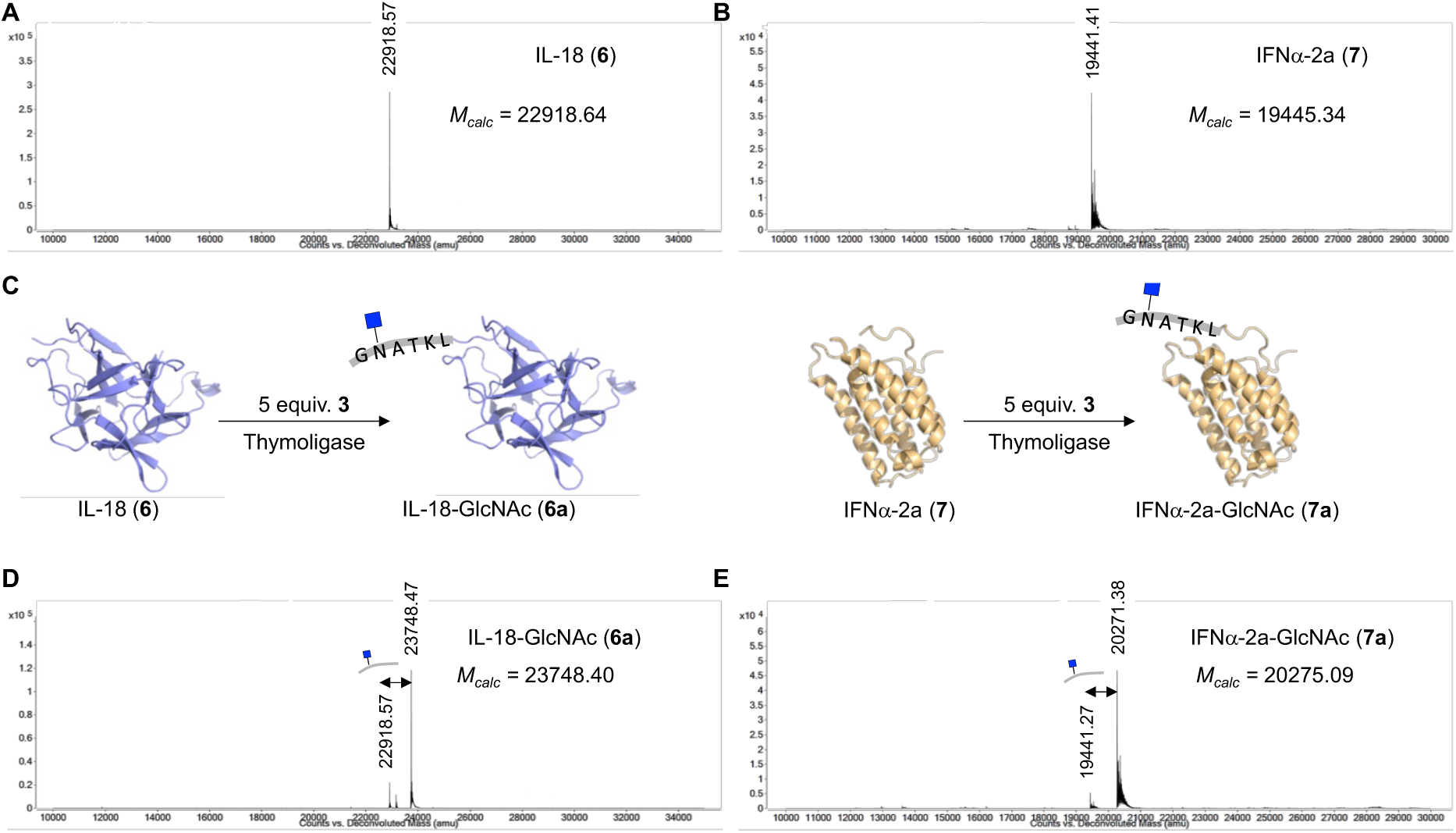
Chemoenzymatic synthesis of IL-18 and IFNα-2a glycovariants **6a** and **7a**. A-B) Deconvoluted mass spectra of IL-18 (**6**) and IFNα-2a (**7**) recombinantly expressed using *E. coli*. C) Reaction of glycopeptide **3** with IL-18 (**6**) or IFNα-2a (**7**) provided homogeneous *N*-acetyl glucosamine modified proteins **6a** and **7a**. D-E) Deconvoluted mass spectra of IL18-GlcNAc (**6a**) and IFNα-2a-GlcNAc (**7a**).

### *N*-Glycosylation of Glyco-tagged Proteins

Since the favorable impact of sialylation on serum half-life of biotherapeutics is well known,^52,53,54^ we focused on modifying GlcNAc-containing IL-18 (**6a**) and IFNα-2a (**7a**) with a complex-type biantennary *N*-sialoglycan. As a convergent approach for *N*-glycoprotein assembly, we explored glycosylation by treatment with endo-glycosidase (ENGase) mutants in combination with sugar oxazolines as activated donor substrates.^55,56^ While wild-type ENGases catalyze the cleavage of the chitobiose core (GlcNAc-β-1,4-GlcNAc) of *N*-glycoproteins, specific ENGase mutants can catalyze the reassembly of custom-made homogeneous *N*-glycan epitopes on the newly exposed GlcNAc site. We selected a mutant ENGase from *Coprinopsis cinerea* (EndoCC1-N180H) ^57^ to modify **6a** and **7a** with oxazoline **8** to give glycoproteins **6b** and **7b**, respectively. This mutant ENGase was selected because it displays a similar specificity as EndoM to transfer biantennary sialoglycans, it is more easily expressed and more thermostable.^58^ Disialoglycan oxazoline **8** was prepared from the corresponding sialoglycopeptide isolated from egg yolk powder^59^ by treatment with EndoS to cleave the glycosidic bond of the chitobiose core followed by reaction with 2-chloro-1,3-dimethylimidazolium (DMC) in the presence of triethylamine to convert the reducing *N*-acetyl-glucosamine moiety into an oxazoline (Figure 4A).^60^ The transglycosylation was carried out using 40 equivalents of **8** and 0.5 mol% EndoCC1-N180H in 50 mM Tris (pH 7.5) at room temperature for 1 h. If starting protein remained, an additional portion of oxazoline **8** was added and incubation was continued for 30 min (Figure 4B). Analysis of the reaction mixture by intact protein MS indicated conversions for IL-18-GlcNAc (**6a**) and IFNα-2a-GlcNAc (**7a**) of ∼90% and ∼51%, respectively. Disialylated IL-18 (IL-18-S2G2, **6b**) was purified by Strep-tag affinity chromatography (Figure 4C for intact MS data). In the case of IFNα-2a, no affinity tag was incorporated in the protein construct and therefore EndoCC1-N180H, which has an His_8_-tag, was removed by Ni-NTA affinity chromatography, and the resulting protein was subjected to Concanavalin A affinity chromatography to afford glycosylated IFNα-2a (**7b**). To further remove incomplete modified products, the protein was subjected to size exclusion chromatography using a ReproSil 125 SEC column, which gave homogeneous **7b** (Figure 4D). These results indicate that the combination of site-selective GlcNAc-peptide ligation and chemoenzymatic glycan elaboration enables the installation of well-defined complex glycans within a target protein. The results demonstrated the feasibility of the glyco-tagging methodology to transform bacterially expressed proteins into well-defined glycosylation variants.

**Figure 4.**
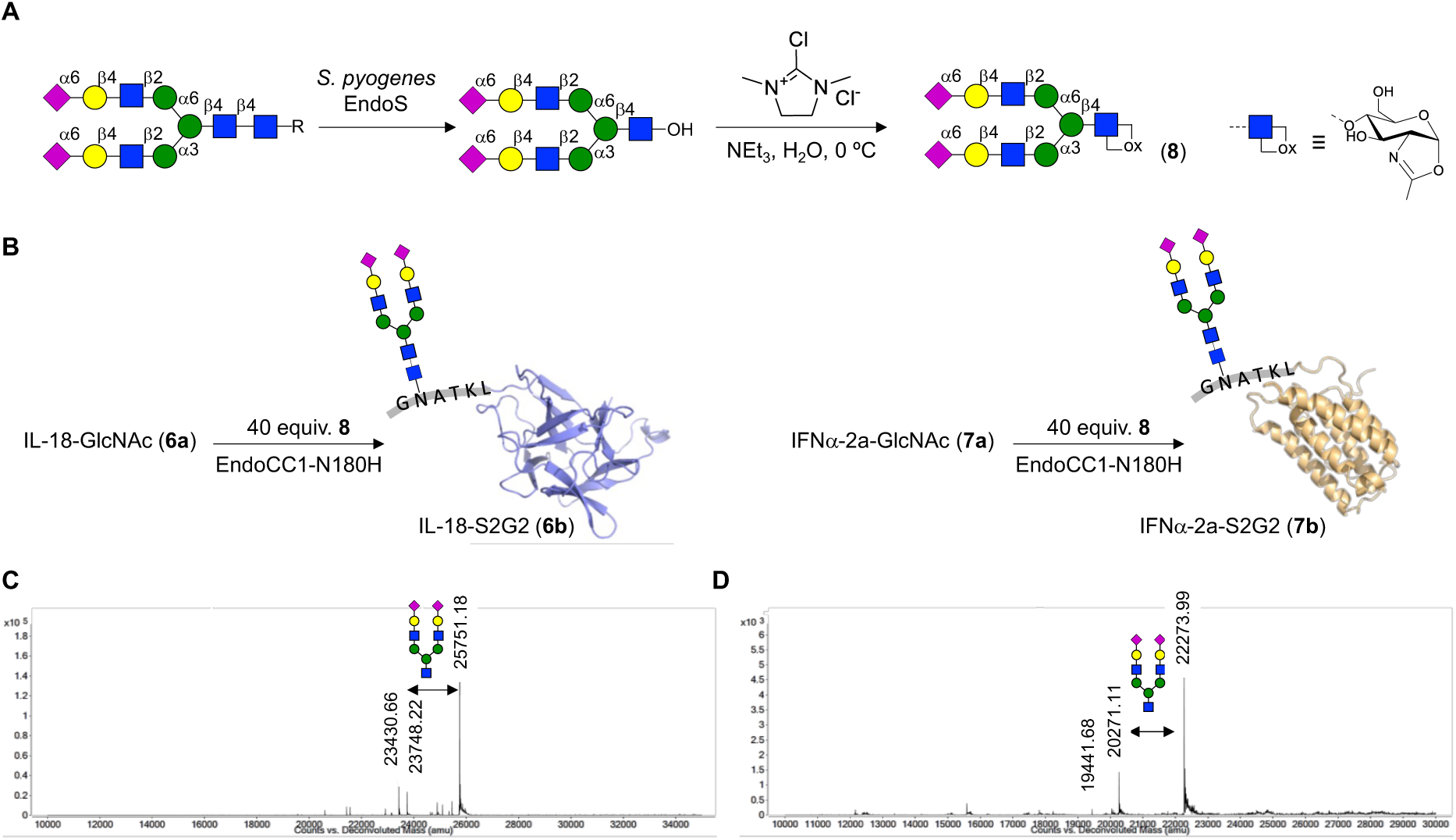
Chemoenzymatic protein remodeling to afford sialylated IL-18 and IFNα-2a glycovariants **6b** and **7b**. A) Conversion of sialoglycopeptide (SGP) isolated from egg yolk powder into oxazoline **8**. B) Protein-glycopeptide conjugates **6a** and **7a** were transformed into complex-type glycosylated variants **6b** and **7b** by reaction with oxazoline **8** in the presence of EndoCC1-N180H. C-D) Deconvoluted mass spectra of glycovariants (**6b**) and (**7b**).

### The Effect of Glycosylation on Biological Activities of Glyco-tagged Cytokines

The influence of protein glyco-tagging on biological properties was evaluated by comparing the pro-inflammatory and anti-proliferative capacities of modified IL-18 and IFNα-2a, respectively, with their unmodified counterparts (Figure 5A-C). IL-18 is a pro-inflammatory cytokine that facilitates type 1 immune responses. In the presence of IL-12, it stimulates IFN-γ production by CD4+ and CD8+ T cells as well as natural killer (NK) cells.^48^ Therefore, IFN-γ release was measured upon the stimulation of human PBMCs with IL-18 and its glyco-tagged variants (0.5 to 50 nM) in the presence of IL-12 (Figure 5A). Stimulation with glyco-tagged variant **6a** and **6b** resulted in comparable IFN-γ release as the non-modified IL-18. To assess whether the natural high-affinity decoy receptor IL-18 BP, a secreted checkpoint inhibitor in cancer therapy,^48^ can interfere with induced IFN-γ release by IL-18 and its glyco-tagged variants, competition binding assays were conducted (Figure 5B). In fact, IFN-γ production stimulated by 10 nM of (glyco-tagged) IL-18 variants was overall diminished in the presence of IL-18 BP (0.5 and 10 nM), ranging from 35-90% at a concentration of 10 nM IL-18 BP. Overall, these data imply that modification by glyco-tagging does not compromise the biological activity of IL-18 variants, compared to its unmodified counterpart.

**Figure 5.**
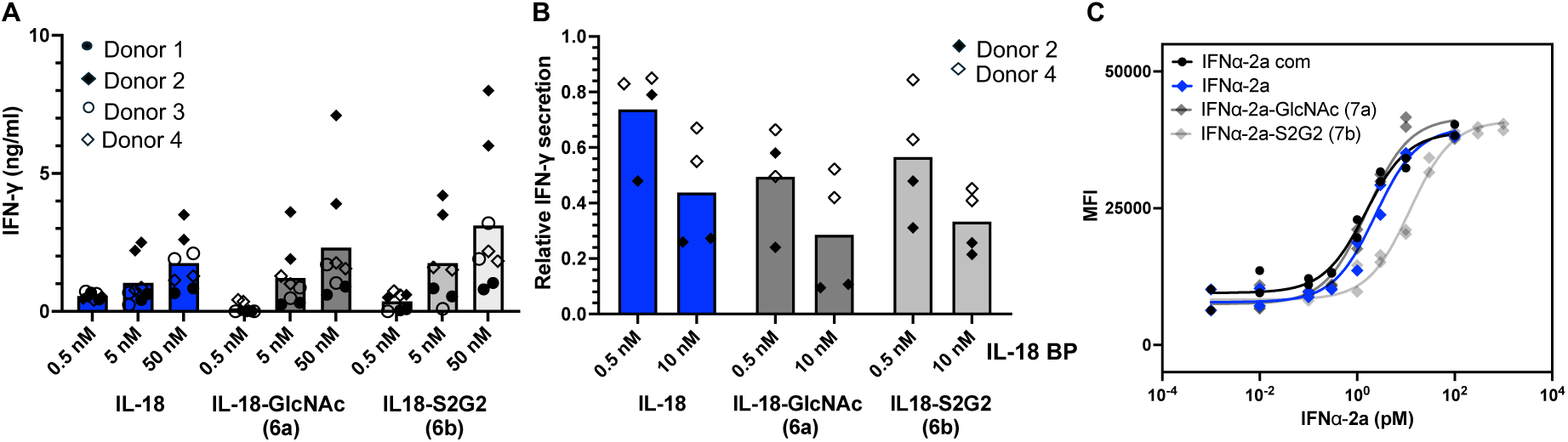
Biological activities of glycan-tagged variants of IL-18 (**6a**, **6b**) and IFNα-2a (**7a**, **7b**). A) Human PBMCs (donors n = 4, biological duplicates) were stimulated with IL-18 variants (0.5, 5.0 and 50 nM) in presence of IL-12 (10 ng/mL) and IFN-γ release was monitored. Data are represented as means combined with each replicate. Inhibition of IL-18 (10 nM) induced IFN-γ release by the decoy protein IL-18 BP (0.5 and 10 nM) is shown as biological replicates (donors n = 2) and is normalized to the average release of each glyco-tagged variant at 10 nM in absence of IL-18 BP. C) CFSE-labelled Daudi cells were stimulated with half-logged concentrations of IFNα-2a variants (1000 to 0.01 pM) in biological duplicates and the mean fluorescence intensity (MFI) was measured after 5 days. EC_50_ values were calculated by sigmoidal curve fitting with a constrained hill slope.

IFNα-2a variants **7a-b** were applied to CFSE (CarboxyFluorescein Succinimidyl Ester)-labelled Daudi cells and the impact of the glyco-tags on its anti-proliferative activity ^49^ was evaluated by tracking mean fluorescence intensities (MFI) - i.e., high MFI implies reduced proliferation and increased anti-proliferative potency (Figure 5C). It was found that purchased and heterologously expressed IFNα-2a (**7**) and the glyco-variant **7a** showed comparable anti-proliferative potency (higher MFI values) while a higher concentration of glyco-variant **7b** was required to achieve the maximal anti-proliferative capacity.

### Chemo-enzymatic Synthesis of Glyco-insulin Variants

Insulin is a pancreatic polypeptide hormone that is used for the treatment of diabetes. Its endogenous form produced by β-cells in response to glucose has a short half-life (∼5 min in human plasma) and is prone to oligomerization, aggregation and fibrillization, which leads to decreased efficacy and potential adverse effects upon its application.^61,62^ Different approaches have been pursued to improve the properties of insulin such as the use of additives and chemical modification. Peglispro, a PEG-modified form of insulin, was developed to provide more stable blood glucose levels over an extended period. However, due to side effects caused by PEG, alternative methods such as glycosylation have been also explored.^63^ Several glyco-insulin variants have been synthesized employing various approaches, although they require demanding chemical synthesis or low yielding conjugation approaches.^13,18^ In view of these earlier examples, we decided to test our tagging methodology for the synthesis of glyco-insulin, using intact insulin from commercial sources as starting material.

Human insulin is composed of an A and B peptide chain (A: amino acids A1–A21, B: amino acids B1–B30) that are connected by two interchain disulfide bonds. The two *N*-terminal amino acids of the A chain are Gly-Ile and that of the B chain Phe-Val, which were both expected to be good substrates for peptidoligases (Figure 6A).^64^ Treatment of native human insulin (**9**) with two equivalents of glycopeptide ester **1** in the presence of omniligase-1 resulted mainly in the formation of mono-ligated insulin as indicated by MS analysis (Figure 6B). Starting material and a small amount of bis-ligated insulin was also detected (56% conversion into monoligated insulin with an estimated selectivity of 86%). The use of a large excess of glycopeptide ester **1** (20 eq., Figure S8 for more details) resulted in a larger conversion but the mono-ligated insulin (INS-GlcNAc) was still the main product albeit with lower selectivity (70% conversion with 66% selectivity for mono-ligated product). On the other hand, full conversion of insulin into bis-ligated insulin **9b** (A1,B1-INS-2xGlcNAc, Figure 6C) was achieved by using thymoligase in the presence of 2 eq. of **1**, after a reaction time of 2 h, as indicated by MS analysis. Thymoligase was developed to accept negatively charged amino acids (Asp) at position P1’^46^ and demonstrated superior activity for protein glyco-tagging compared with omniligase-1.

**Figure 6.**
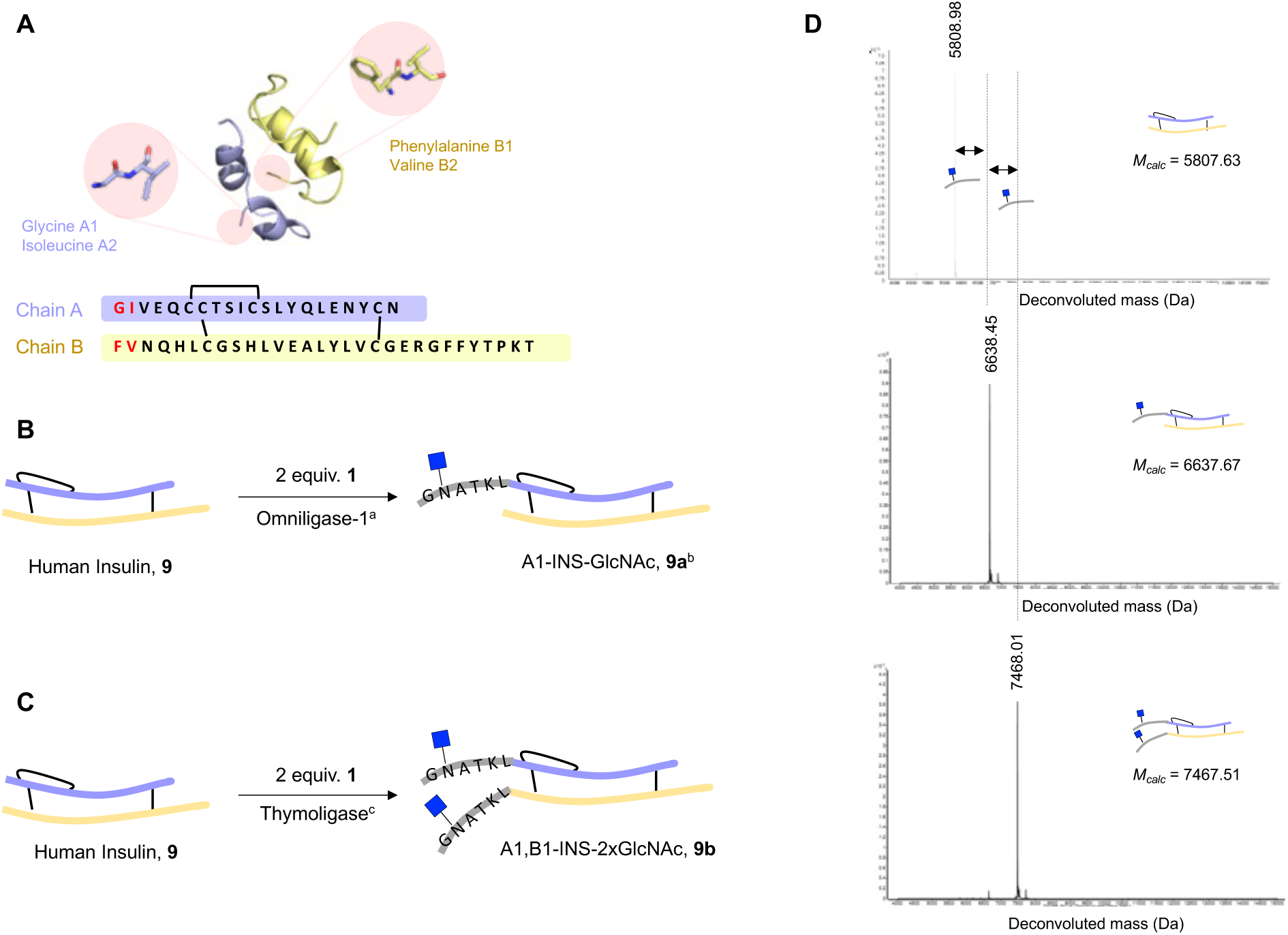
Peptidoligase-catalyzed human insulin-glycopeptide conjugation. A) Structure (pdb file 1a7f) and amino acid sequence of native human insulin. Amino acids pairs at both chain A and B *N*-terminus are highlighted in red. B) and C) Chemoenzymatic synthesis of *N*-GlcNAc insulins **9a** and **9b**. Conversions were estimated by mass spectrometry analysis. ^a^ Reaction conditions were 200 µM **9**, 400 µM 3, 1.5 mol% omniligase-1, 100 mM tricine (pH 8.0), RT, 2 h. 56% of the starting material was converted into monoligated insulin. ^b^ **9a** was obtained in 30% isolated yield. Selectivity of the reaction was determined by treatment of the sample with DTT and detection of the individual peptides. ^c^ Reaction conditions were 200 µM **9**, 400 µM **3**, 1.5 mol% thymoligase, 100 mM tricine (pH 8.0), RT 2 h. If unmodified protein or monoligated insulin remained after 2 h a second portion of glycopeptide **3** was added and allowed to react for additional 2 h. Addition of two units of glycopeptide occurred in >95% conversion. **9b** was obtained in 40% isolated yield. D) Deconvoluted mass spectra of intact human insulin **9** and glycoinsulin variants **9a**-**b**.

The mono- and bis-modified glyco-insulins were purified by preparative reverse phase HPLC using a C18 column and their identity and structural integrity analyzed by ESI-QTOF mass spectrometry. Deconvoluted mass spectra of **9a** and **9b** displayed single peaks with m/z of 6637 Da and 7468 Da, respectively, corresponding to the addition of one and two GlcNAc-containing peptides (Figure 6D). The m/z peaks corresponding to individual A and/or B chains were not detected, supporting the compatibility of the methodology with the presence of disulfide bonds. The precise identity of mono-ligated insulin, compound **9a**, was ascertained by treatment with dithiothreitol (DTT) followed by MS analysis of the individual peptides. m/z signals corresponding to glycopeptide addition to the A-chain and almost no modified B-chain were detected (Figure S9 for additional details) and the selectivity appears to be greater than >95%. These results indicate that the substrate specificity of omniligase-1 resembles that of its parent peptiligase, which prefers small amino acids in the S1’ pocket such as Gly, Ala or Ser.^39,40^ In the case of thymoligase, S1’ pocket was engineered to introduce a L217R point mutation that could promote favorable cation-π interactions. It is possible that this modification also facilitates the recognition of Phe at the *N*-terminus of the B-chain of insulin, thus providing full conversion to bis-ligated product **9b**.

Next, attention was focused on the modification of GlcNAc moieties of **9a** and **9b** with a complex sialylated *N*-glycan (Figure 7A-B). Thus, treatment of **9a** and **9b** with oxazoline **8** in the presence of EndoCC-N180H for 1 h resulted in the installation of complex-type glycans at the GlcNAc sites. Peaks with m/z of 8640.88 Da and 11473.46 Da matched the expected molecular weights for the addition of one or two complex glycans to afford **9c** and **9d,** respectively (Figure 7C). Also, minor peaks resulting from the loss of sialic acid and the formation of TFA adducts were detected, which most likely arise from in-source MS fragmentation or ion addition.^65^ The glyco-insulins were purified by HPLC using a C18 column to give homogeneous **9c** and **9d** in isolated yields of 50% and 74%, respectively. Thus, the two-step modification offers a highly convergent strategy to prepare well-defined glycosylated insulin variants from readily available native human insulin.

**Figure 7.**
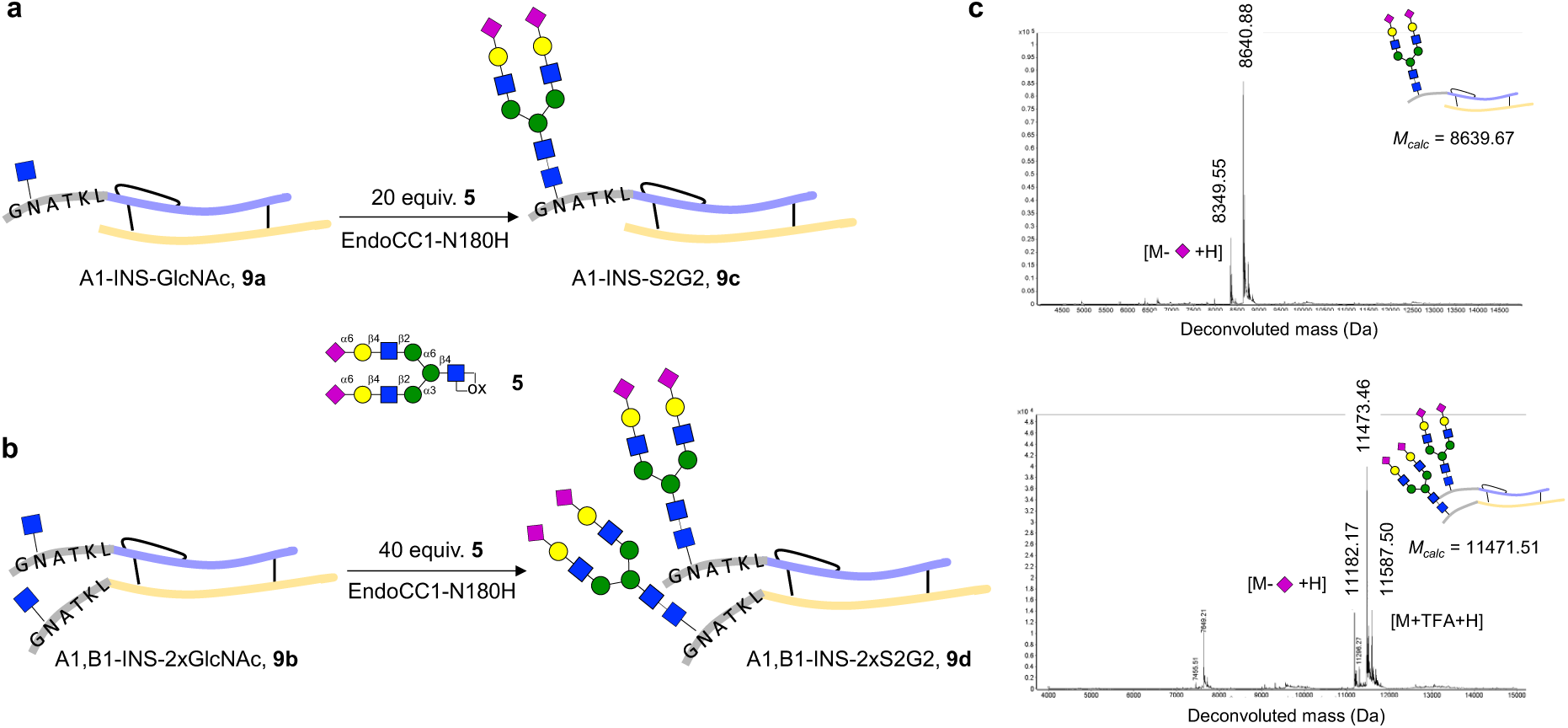
Chemoenzymatic synthesis of disialo-glycoinsulin analogues. A) and B) Chemoenzymatic installation of sialo complex-type biantennary oxazoline **8** into glucosylated insulins **9a** and **9b**. 200 µM **9a/9b**, 20-40 equivalents of **8**, 0.1 mol% EndoCC1-N180H, Tris-HCl 20 mM (pH 7.0), RT, 1 h. If starting protein remained a second portion of oxazoline **8** was added and allowed to react for an additional 1 h. Glycoinsulins **9c** and **9d** were obtained in 50% and 74% isolated yield, respectively. C) Deconvoluted mass spectra of glycoinsulin variants **9c-d**.

### Physical Properties of Glyco-insulin Derivatives

In the pancreas, human insulin is stored in β-cells as an inactive and symmetric Zn^2+^ coordinated hexamer. In response to an increase in blood glucose levels, insulin is released to the blood stream where it rapidly dissociates to form physiologically active monomers. While the inactive hexamer is rather stable, the monomeric form is not and can partially unfold and then self-associate into oligomers, which are considered hallmarks of the pre-fibrillar phase that rapidly evolve to form larger aggregates and fibrils. Such aggregation compromises the therapeutic use of insulin.^66^

Diffusion ordered NMR spectroscopy (DOSY) can provide information about the size/shape of molecules.^67^ Therefore, we used this experimental approach to investigate the oligomeric state of the glyco-insulin variants in solution.^68^ First, the insulin derivatives were investigated at low pH and low salt concentration because these conditions are known to greatly reduce aggregation. Thus, 50 µM buffered solutions of human insulin **9** and the different glyco-insulins **9a-d** at pH 1.6 were prepared and 2D ^1^H-DOSY-NMR experiments were acquired to provide diffusion coefficients, D (Figure 8A). All DOSY spectra displayed a single set of NMR signals (Figure S14), indicating that human insulin **9** and its glycovariants **9a-d** are present in a single oligomeric state or if they exist as an ensemble of different oligomeric forms, they are in fast exchange.

**Figure 8.**
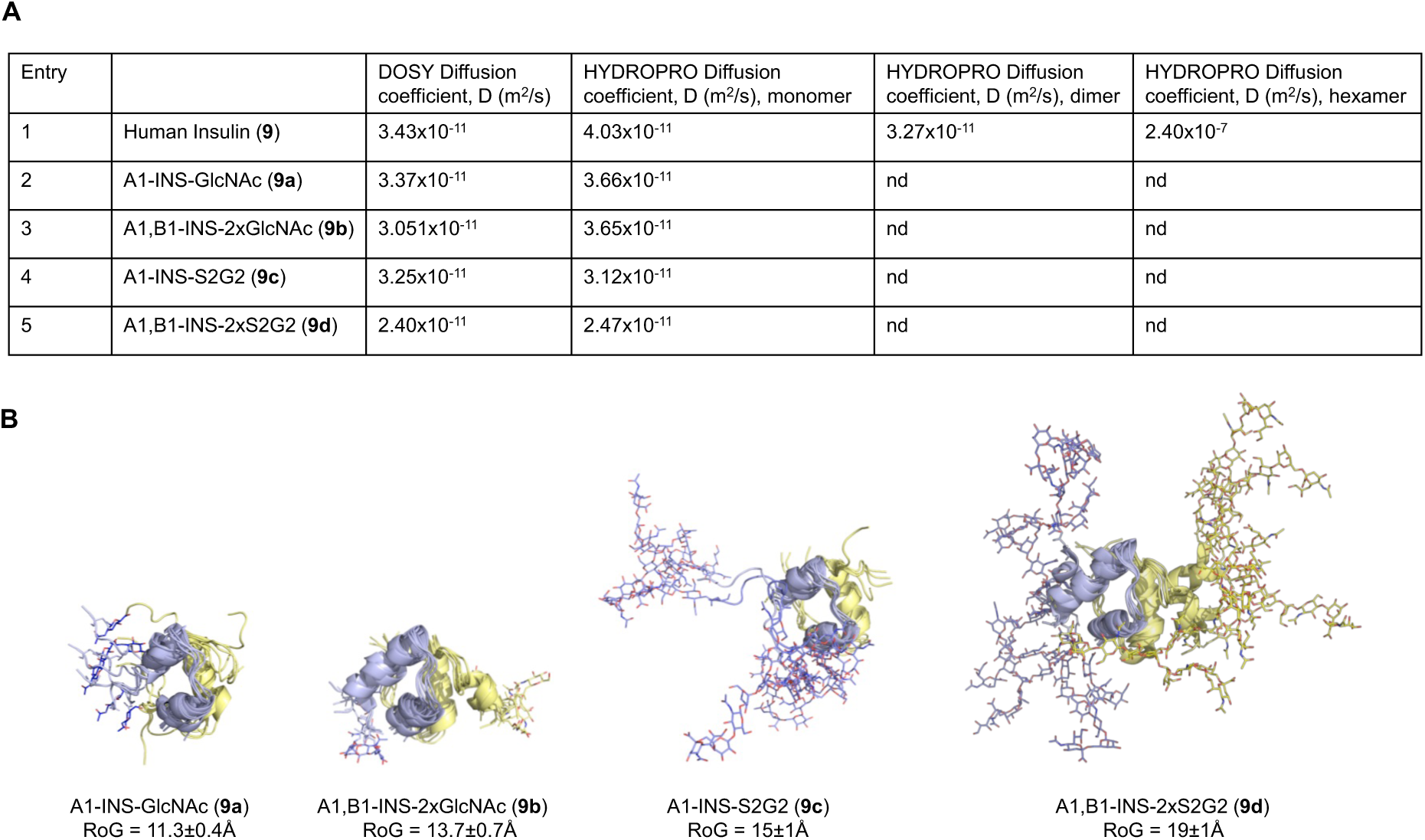
Analysis of the oligomeric state of human insulin **9** and derived glycoinsulins **9a-d**. A) Diffusion coefficients of human insulin **9** and glycoinsulin variants **9a-9d** determined by diffusion ordered spectroscopy NMR experiments (2D ^1^H-DOSY-NMR). Calculated diffusion coefficients of human insulin monomer, dimer and hexamer as well as those for glycoinsulins **9a**-**9d** monomers were also indicated. B) 3D structural models of glycoinsulins **9a**-**9d**. The superimposition of several MD snapshots showed the increased sampled conformational space of disialoglycosylated insulins and the impact on the radius of gyration of the molecule.

To discern the oligomeric state of human insulin under the employed experimental conditions, theoretical diffusion coefficients were calculated using the simulation software HYDROPRO and compared with the experimental values. X-ray crystal structures of human insulin monomer, dimer, and hexamer (PDB ID 3aiy) were used as coordinate input files. The experimental diffusion coefficient of bovine pancreatic ribonuclease A, a small globular protein of 13.7 kDa (PDB ID 3rn3), was used as reference to standardize the computed values. In agreement with previously reported values, the determined diffusion coefficient for unmodified human insulin **9** matched a dimer as the main species (Figure 8A, entry 1).^69^ In particular, 4:1 ratio in favor of insulin dimer would account for the measured diffusion coefficient, when monomer and dimer species were considered to calculate the weighted mean value. For comparison purposes, 3D structural models of **9a**-**9d** monomers were built using AlphaFold in combination with the glycoprotein builder module implemented in GLYCAM-Web portal and their dynamic behavior was interrogated by molecular dynamics (MD) simulations (Figure 8B). Installation of GlcNAc or S2G2-bearing glycopeptides increased the radius of gyration (RoG) and therefore the hydrodynamic radius of the analyzed proteins. Predicted RoG and diffusion coefficients for disialoglycoinsulins **9c** and **9d**, carrying highly hydrophilic and conformationally flexible *N*-glycans, were substantially higher. Interestingly, experimentally determined diffusion coefficients for **9a**-**9d** were in close agreement to the calculated values of the monomeric species (Figure 8A, entries 2-5). Thus, based on their diffusion properties, peptide glyco-tagging and especially the introduction of large *N*-glycans, such as in the case of glycoinsulins **9c** and **9d**, shifts the oligomeric equilibrium of human insulin towards monomers.

To investigate the amyloidogenic properties of the glyco-tagged insulin variants, experimental conditions that favor aggregation were applied.^70^ Thus, 50 µM protein samples in buffered solutions at pH 1.6 and containing 150 mM NaCl were incubated at 60 °C and their aggregation kinetics were analyzed by NMR. In the case of human insulin **9**, acquisition of ^1^H-NMR experiments allowed detection of the decay of the ^1^H signal intensities over time. After 12 h of incubation, the signal intensity was reduced to 2% (Figure 9A-B). Close monitoring allowed identifying a sigmoidal kinetics, which comprised a typical lag phase followed by a faster growth/aggregation step. No new peaks appeared. These observations are indicative of the conversion of small size oligomers (NMR visible species) into large size oligomeric species whose NMR signals are broadened and not NMR-visible anymore. Remarkably, almost no aggregation of glycoinsulin variants **9a**-**9d** was observed, and the decay in signal intensity was less than 10% after long incubation of 49 h (for A1-INS-S2G2, **9c**) (Figure 9A-B).

**Figure 9.**
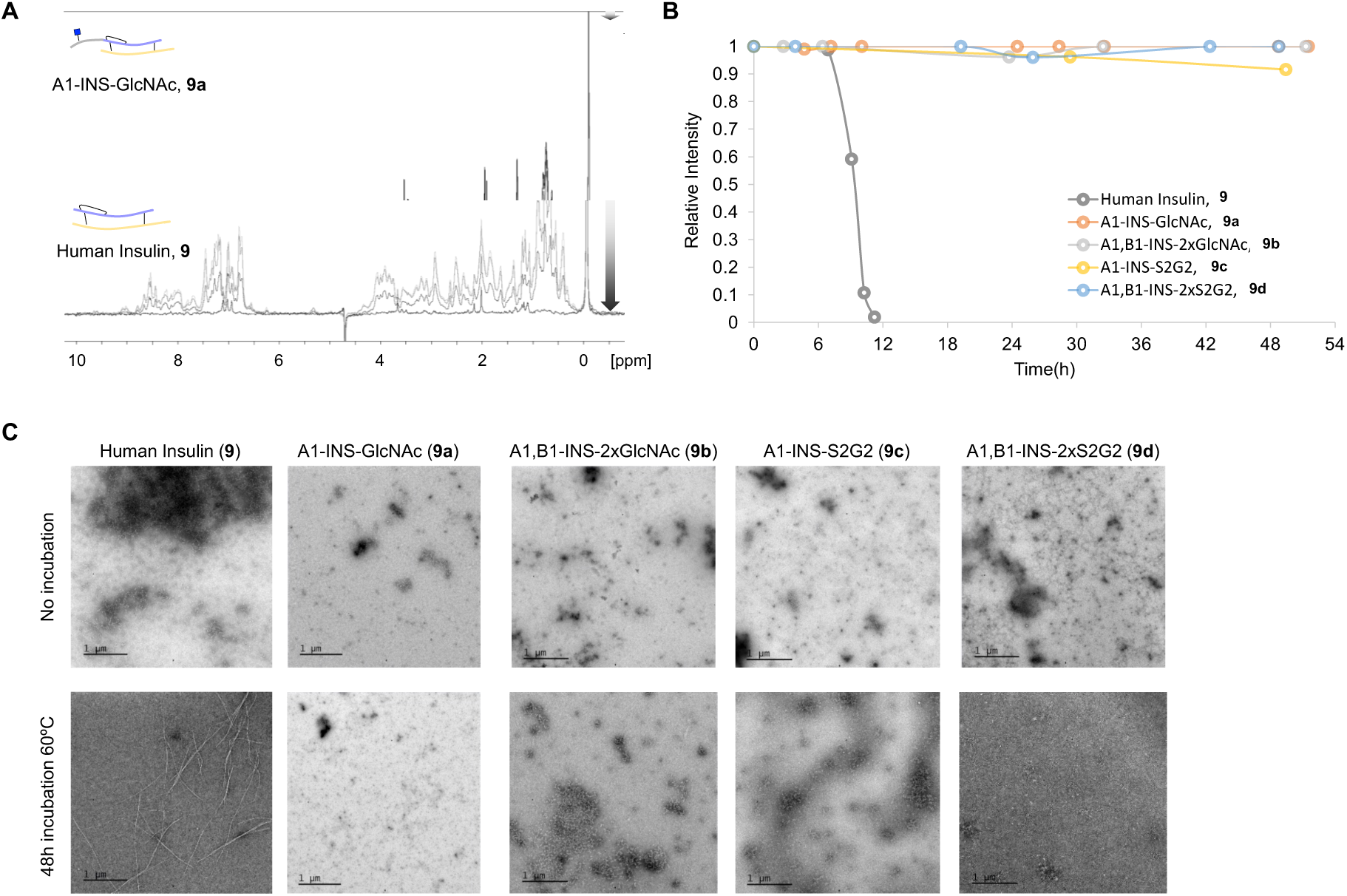
Aggregation properties of human insulin 9 and glycoinsulin variants **9a-d**. A) ^1^H-NMR monitoring of signal intensity changes for human insulin **9** and glycosylated insulin **9a** over sample incubation at 60 °C. A similar trend was observed for synthesized insulins **9b**-**9d**. B) Decay of relative intensity of protein signals over time. 50 µM samples in phosphate buffer pH 1.6 containing 150 mM NaCl were analyzed. C) TEM images of 30 µM insulin **9** and glycoinsulin **9a**-**d** samples before and after 48 h incubation at 60 °C.

Transmission electron microscopy (TEM) was employed to further analyze possible aggregation of insulin and insulin derivatives after incubation period of 48 h at 60 °C. As expected, only fibril formation was observed for human insulin **9**, while large structures were absent for the glycoinsulins **9a**-**d** (Figure 9C). Collectively, the results demonstrate that glycosylation protects insulin from aggregation. Although the mechanism underlying insulin aggregation and fibrillization is not fully understood, it is widely accepted that hydrophobic interactions facilitate the unfolding and misfolding of the polypeptide chains resulting in agglomeration, fibrillogenesis, and precipitation.^71,72^ The introduction of hydrophilic entities such as glycans, may disrupt unfavorable hydrophobic interactions.

### Biological Activities of Glycoinsulin Variants

Binding of insulin to its cell surface receptor on metabolic cells triggers a complex cascade of intracellular events, inducing amongst other events, glucose uptake by the translocation of Glucose transporter type 4 (GLUT4).^73^ A central event of this signaling cascade is the phosphorylation of protein kinase B (PKB), on its Thr308 residue (by PDK1) and Ser473 (by mTORC2), resulting in kinase activation and subsequent phosphorylation of multiple downstream targets, such as PRAS40, an inhibitory binding partner for mammalian target of rapamycin (mTOR).^74^ Activated mTOR regulates upstream a plethora of cell-specific biological processes such as glucose metabolism, lysosomal biogenesis, and lipid synthesis. To assess the impact of insulin glycosylation on downstream signaling, fully differentiated Simpson-Golabi-Behmel Syndrome (SGBS) human adipocyte cells were stimulated with unmodified and the glyco-tagged insulin variants **9a-d** at 5 and 100 nM for 5 min. Thr308- and Ser473-PKB phosphorylation as well as phosphorylation of Thr246-PRAS40 substrate were quantified by Western Blotting (Figures 10A and S16A). The sensitivity of the test system was assessed by the application of all variants at a concentration of 5 nM. All variants were able to induce phosphorylation at Thr308- and Ser473-PKB, however, different levels of phosphorylation were observed for the various glyco-variants in particular at a low concentration of 5 nM (Figure 10B). Introduction of single GlcNAc moieties at the A-chain or both chains (**9a** and **9b,** respectively) had only a marginal effect on the phosphorylation levels when compared to unmodified insulin. In contrast, the installation of large complex glycans, especially in the case of variant **9d** resulted in reduced phosphorylation. Further downstream, a similar behavior was observed by quantifying the phosphorylation levels of the mTOR inhibitory substrate PRAS40 (Figure 10C) and the phosphorylation of various other PKB substrates using an antibody recognizing its consensus motif for phosphorylation (Figure S16B).

**Figure 10.**
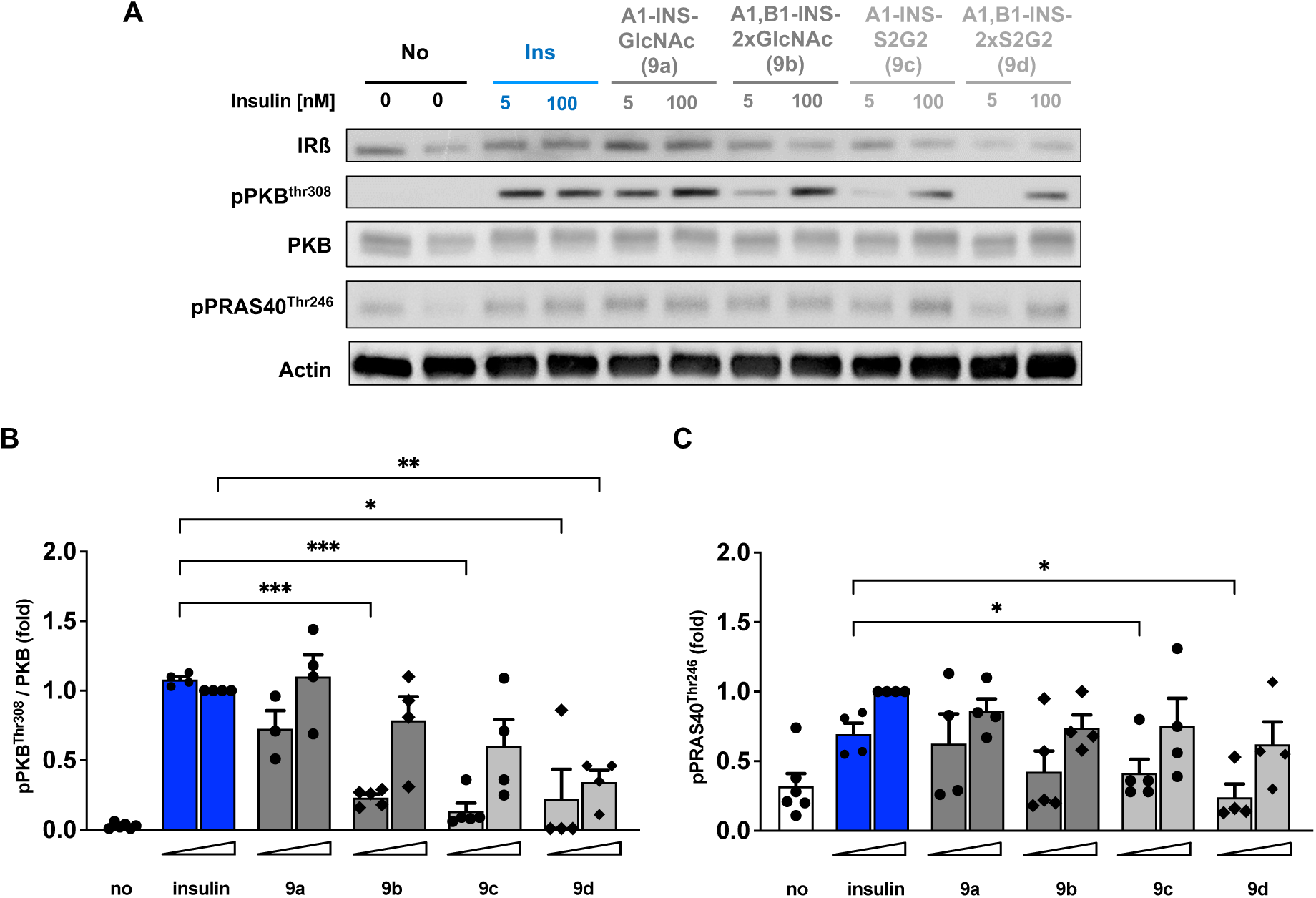
Insulin signaling by human insulin and glycoinsulin variants **9a-d**. SGBS human adipocyte cells (n=4) were differentiated for 14 days in adipogenic medium and serum-starved for 16 h before stimulation for 5 min. with human insulin and the glycovariants **9a**-**d** at 5 and 100 nM. The cells were lysed and phosphorylation of PKB at residue Thr308 and PRAS at residue Thr246 was detected by Western Blot. Phosphorylation levels were quantified for the phosphorylation sites B) Thr308 of PKB and C) Thr246 of the substrate PRAS40 and normalized against the non-phosphorylated protein. Data are represented as means *+/-* the standard error of the mean (SEM). Statistical significance was tested with a mixed effect model using Restricted Maximum Likelihood (REML) for data fitting (implemented in Prism 10).

## CONCLUSIONS

A chemoenzymatic methodology for glyco-tagging of native proteins is described that exploits peptidoligases to fuse a peptide ester carrying an *N*-acetyl-glucosamine (GlcNAc) moiety to the *N*-terminus of a protein generating a *N*-GlcNAc modified glycoprotein. The GlcNAc moiety of the resulting glycoprotein can be elaborated into complex glycans by *trans*-glycosylation using well-defined sugar oxazolines and mutant forms of endo β-*N*-acetylglucosaminidases (ENGases). The workflow is orthogonal with high yielding protein expression in *E. coli* that cannot glycosylate proteins. The strategy was employed for the preparation of glycosylation variants of several proteins, including the cytokines, IL-18 and IFNα-2a, and human insulin.

The glycovariants maintained protein functionality, while in the case of human insulin the new constructs displayed improved physicochemical properties. Engineered mutants of peptidoligases have been described that have different activities for the *N*-terminal sequences,^64^ which offers the prospect of modifying many different native proteins by glycotags. Additionally, chemoenzymatic methodologies have been described that make it possible to prepare large panels of *N*-linked glycans starting from a readily available bi-antennary glycopeptide isolated from egg yolk powder that can be converted into multi-antennary *N*-glycans by using recombinant glycosyl transferases and modified sugar nucleotide donors.^75^ The combination of these methodologies paves the way to modify native proteins in a well-defined manner with a diverse range of glycans for structure-function *in-vitro* and *in-vivo* studies and it is to be expected to provide a much needed tool for improving properties of therapeutic proteins.

## ASSOCIATED CONTENT

### Supporting Information

The Supporting Information is available with:

Methods, analytical data, and additional figures (PDF).

## AUTHOR INFORMATION

### Authors

**Ana Gimeno** - Chemical Biology and Drug Discovery, Utrecht Institute for Pharmaceutical Sciences, Utrecht University, Utrecht 3584 CG, The Netherlands. Present address: CICbioGUNE, Basque Research & Technology Alliance (BRTA), Bizkaia Technology Park, Building 800, 48160 Derio, Bizkaia, Spain; Ikerbasque, Basque Foundation for Science, Plaza Euskadi 5, 48009 Bilbao, Spain; orcid.org/0000-0001-9668-2605

**Anna M. Ehlers** - Chemical Biology and Drug Discovery, Utrecht Institute for Pharmaceutical Sciences, Utrecht University, Utrecht 3584 CG, The Netherlands.

**Sandra Delgado** - CIC bioGUNE, Basque Research & Technology Alliance (BRTA), Bizkaia Technology Park, Building 800, 48160 Derio, Bizkaia, Spain.

**Jan-Willem H. Langenbach** - Chemical Biology and Drug Discovery, Utrecht Institute for Pharmaceutical Sciences, Utrecht University, Utrecht 3584 CG, The Netherlands.

**Leendert J. van den Bos** - EnzyTag BV, Daelderweg 9, NL-6361 HK Nuth, The Netherlands; orcid.org/0000-0003-2410-5838

**John A.W. Kruijtzer** - Chemical Biology and Drug Discovery, Utrecht Institute for Pharmaceutical Sciences, Utrecht University, Utrecht 3584 CG, The Netherlands.

**Bruno G.A. Guigas** - Leiden University Center of Infectious Diseases, Leiden University Medical Center, Leiden 2333 ZA, The Netherlands

### Author Contributions

The studies were conceived by G.J.B. and A.G. G.J.B. was responsible for overall project management. A.G. performed ligations, glyco-remodelling, compound characterization, NMR and computations studies of insulin derivatives. A.M.E. performed biological experiments.

L.J.v.d.B. provided reagents and conditions for ligations. S.D. performed TEM studies.

J.A.W.K developed approach for peptide ester synthesis. J.W.H.L prepared oxazolidines.

B.G.A.G. supervised biological evaluation of glycoinsulin variants. A.G., A.M.E., and G.J.B. wrote the paper. All authors have given approval to the final version of the manuscript.

### Notes

L.J.v.d.B. is a minority shareholder and employee of EnzyTag. The other authors declare no competing financial interest.

## ACKNOWLEDGMENTS

This research was supported by a grant from NWO TA PPS fund (ENPPS.TA.019.008 to G.J.B.). We thank Frank Otto for technical help with SGBS cells, Dr. Anna Koijen (EnzyTag) for discussions on peptidoligase technology and providing omniligase-1 and thymoligase.

