## Supplementary material for "A Chemoenzymatic Strategy for Site-Specific Glyco-Tagging of Native Proteins for the Development of Biologicals": SI

**TABLE OF CONTENTS**


1. **Materials** **S2**
2. **Methods** **S2**
   - **2.1. Chemical and Chemoenzymatic Synthesis** **S2**
   - **2.2. Protein Expression and Purification** **S12**
   - **2.3. Chemoenzymatic Protein Modification** **S15**
   - **2.4. Structural and Biophysical Analysis of Insulin Variants** **S24**
   - **2.5. Biological Assays of IL-18 and IFNa2a Variants** **S26**
   - **2.6. Biological Assays of Insulin Variants** **S27**
3. **References** **S30**

### 1. Materials

*Proteins.* Bovine serum albumin and human insulin were purchased from Sigma-Aldrich and used without further purification.

*Peptiligases.* Omniligase-1 and thymoligase were kindly provided by Enzytag.

*Glycosidases and glycosyltransferases.* α-(2,6)-sialyltransferase (ST6Gal1),[1] endoglycosidase wild-type from *Streptococcus pyogenes* (Endo S)[2] and endoglycosidase mutant N180H from *Coprinopsis cinerea* (EndoCC1-N180H)[3,4] were prepared as reported. Calf intestine alkaline phosphatase (CIAP) and Neuraminidase from *Clostridium perfringens* were purchased from Invitrogen™ [Cat# 18009027] and New England Biolabs [Cat# P0720L], respectively.

### 2. Methods

#### 2.1. Chemical and Chemo-enzymatic Synthesis of Small Molecules

##### 2.1.1. General experimental procedures

Unless otherwise stated, all reagents were of synthetic grade and used as received. Dichloromethane, and *N*, *N*-dimethylformamide used for synthesis were anhydrous grade and obtained from a solvent purifier (MBraun SPS 800). Organic solvents for work-up procedures were technical grade and obtained from VWR Chemicals. Deuterated solvents for NMR experiments were obtained from Cambridge Isotope Laboratories. All moisture sensitive reactions were performed under an argon atmosphere and in the presence of molecular sieves. Unless otherwise stated, all reactions were carried out in glassware material with magnetic stirring. Molecular sieves were flame-dried in vacuo immediately prior to use. TLC analysis was performed using precoated silica gel 60 F-254 plates (Merck) with detection by UV (254 nm) and when applicable by spraying with 20% sulfuric acid in EtOH followed by charring at ~150 °C. Flash column chromatography was performed on silica gel G60 (Silicycle, 60-200µm, 60Å). $^{1}$H and $^{13}$C NMR spectra were recorded on a 400 MHz Varian or 600 MHz Brucker NMR spectrometers in $CDCl_3$, $D_2O$, DMSO-$d_6$. Chemical shifts (δ) are given in ppm relative to the residual signal of the deuterated solvent. Coupling constants (*J*) are given in Hz. All $^{13}$C spectra are proton decoupled. $^{1}$H NMR resonances were assigned through standard TOCSY, NOESY and HSQC experiments.

#### 2.1.2. Synthesis of non-canonical amino acids

Fmoc-LeuOCamOH[5] (**1**) and Fmoc-Asn(GlcNAc)[6] (**2**) were prepared using described procedures. The identity of the prepared compounds was confirmed through $^1$H-NMR analysis and comparison with previously reported data.

**2-(((((9H-fluoren-9-yl)methoxy)carbonyl)-L-leucyl)oxy)acetic acid (4).** In a microwave compatible vial (20 mL), Fmoc-Leu-OH (1.77 g, 5.0 mmol) was dissolved in DMF (5 mL). A solution of potassium iodide (0.52 g, 3.0 mmol) and diisopropylamine (1.75 mL, 12.45 mmol) in DMF (25 mL total volume) was prepared. The resulting solution (10 mL) was added to the Fmoc-Leu-OH solution. Finally, tert-butyl bromoacetate (1.46 g, 7.5 mmol) was added and the vial was capped. The reaction mixture was heated to 90 °C (MW) for 10 min. The resulting orange-brown solution was transferred to a separatory funnel. Ethyl acetate (150 mL) was added, and the solution was washed with water (50 mL), 1 M sodium thiosulphate (50 mL), 1M potassium bisulphate (2 x 50 mL), saturated sodium bicarbonate (2 x 50 mL) and brine (50 mL). The combined organic layers were dried over anhydrous $Na_2SO_4$, filtered, and the filtrate was concentrated in vacuo. The crude material was purified by silica gel column chromatography using EtOAc/Hex (2:3 v/v) to afford Fmoc-Leu-OCam-OtBu. The product was directly dissolved in DCM/TFA (1:1 v/v, 80 mL) and the reaction mixture was stirred for 2 h at room temperature (RT). The reaction mixture was concentrated in vacuo and the crude material was purified by silica gel column chromatography using EtOAc/Hex (2:3 v/v). Yield: 79% (over two steps), white foam. $^1$H NMR (400 MHz, CDCl3) δ 7.75 (d, 2H, $J^3_{HH}$ = 7.6 Hz, Ar-H, Fmoc), 7.57 (dd, 2H, $J^3_{HH}$ = 7.7, $J^4_{HH}$ = 3.8 Hz, Ar-H, Fmoc), 7.45 – 7.33 (m, 2H, Ar-H, Fmoc), 7.29 (t, 2H, $J^3_{HH}$ = 7.6 Hz, Ar-H, Fmoc), 5.12 (d, 1H, $J^3_{HH}$ = 8.5 Hz, NH), 4.77 (d, 1H, $J^2_{HH}$ = 16.3 Hz, $CH_2$, OCam), 4.64 (d, 1H, $J^2_{HH}$ = 16.3 Hz, $CH_2$, OCam), 4.42 (m, 3H, $CH_2$, Fmoc + CHα), 4.21 (t, 1H, $J^3_{HH}$ = 6.8 Hz, CH, Fmoc), 1.78 – 1.67 (m, 1H, $CH_2$, Leu), 1.58 (m, 1H, $CH_2$, Leu), 0.95 (d, 6H, $J^3_{HH}$ = 5.5 Hz, $CH_3$, Leu; m, 1H, CH, Leu).

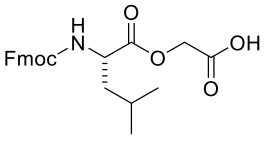


***N*2-(((9H-fluoren-9-yl)methoxy)carbonyl)-*N*4-((2R,3R,4R,5S,6R)-3-acetamido-4,5-dihydroxy-6-(hydroxymethyl)tetrahydro-2H-pyran-2-yl)-L-asparagine (5).** *N*-acetyl-glucosamine (10.0 g, 45.2 mmol) and $NH_4HCO_3$ (100 g, 1.25 mol) were dissolved in water (350 mL) and the resulting cloudy solution was stirred vigorously at RT for 14 days. The solution was concentrated in vacuo and the residue was re-dissolved in water (40 mL) and subsequently

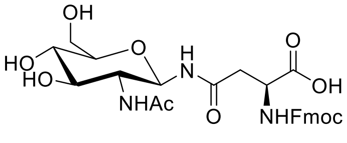

lyophilized, repeating the action three times to remove ammonium bicarbonate. This procedure afforded a white solid powder containing 8:2 mixture of *N*-acetyl-amino glucopyranosyl amine and the starting material as determined by $^{1}$H NMR. The mixture was used without further purification. Therefore, Fmoc-Asp-OtBu (2.59 g, 6.2 mmol), HOBt (0.82 g, 5.78 mmol) and HBTU (6.64 g, 15.77 mmol) were dissolved in DMF (17 mL). A solution of the amine (1.16 g, 4.20 mmol) in DMF (17 mL) was prepared. Both solutions were combined, DIPEA (1.10 mL, 12.6 mmol) was added, and the reaction mixture was stirred at RT for 2 h. The reaction mixture was concentrated in vacuo and the crude product was purified by flash chromatography (3:1, v/v, DCM/MeOH) to afford the corresponding coupling product as a white powder which properties were consistent with previously reported data.[6] The resulting compound (1.468 g, 2.39 mmol) was dissolved in DCM/TFA (20 mL, 50:50 v/v) and the solution was stirred for 3 h at RT. The reaction mixture was concentrated in vacuo, and the resulting syrup was azeotroped with toluene (5 x 20 mL), DCM (3 x 20 mL) and diethyl ether (1 x 50 mL). The residue was dried under reduced pressure to afford compound **2**. Yield: 37% (over 3 steps), amorphous white solid. 1H NMR analysis was consistent with previously reported data [3,4]: (400 MHz, DMSO-d6/$D_2O$) δ 7.82 (d, 2H, $J^3_{HH}$ = 7.5 Hz, Ar-H, Fmoc), 7.63 (d, 2H, $J^3_{HH}$ = 7.5 Hz, Ar-H, Fmoc), 7.36 (t, 2H, $J^3_{HH}$ = 7.4 Hz, Ar-H, Fmoc), 7.26 (t, 2H, $J^3_{HH}$ =7.8Hz, Ar-H, Fmoc), 4.75 (d, 1H, $J^3_{HH}$ = 9.6 Hz, H1, GlcNAc), 4.28 (dd, 1H, $J^3_{HH}$ = 7.6, 5.2 Hz, CH, Asn), 4.23 – 4.10 (m, 3H, $CH_2$, CH, Fmoc), 3.57 (m, 1H, H6α, GlcNAc), 3.49 – 3.28 (m, 3H, H2, H6β, H3, GlcNAc), 3.12 – 3.01 (m, 2H, H4, H5, GlcNAc), 1.72 (s, 3H, NHAc, GlcNAc).

#### 2.1.3. Solid phase peptide synthesis - general procedure

Peptides were synthesized using automated Fmoc solid phase synthesis on a HT12 Liberty Blue™ automated microwave peptide synthesizer using Tentagel™ S-RAM rink amide resin from Rapp polymere (Germany), and DMF as solvent on a 0.2-0.5 mmol scale. The following side chain protecting groups were used: Asn(Trt), Gln(Trt), Glu(OtBu), His(Trt), Lys(Boc), Ser(tBu), Thr(tBu), Trp(Boc),Tyr(tBu). Single coupling reactions were performed using five equivalents of the appropriate Fmoc amino acid, and diisopropylcarbodiimide (DIC), and Oxyma as activator and base, respectively. 20% (v/v) piperidine in DMF was used as a deprotection cocktail. Same conditions were used for the coupling and deprotection of non-canonical amino acids Fmoc-Asn(GlcNAc) and Fmoc-LeuOCamOH. Final capping was performed with 10% (v/v) acetic anhydride in DMF. Peptides were cleaved and side chains were deprotected by incubating the resin with 95:2.5:2.5 ratio of trifluoroacetic acid (TFA), water, and triisopropylsilane (TIPS). The resulting mixture was filtered, and the solution was concentrated to 5 mL on a rotary evaporator. The peptides were precipitated by addition of 10 volumes of diethyl ether:hexanes 1:1, washed twice diethyl ether:hexanes 1:1 and finally with diethyl ether. Peptides were purified by C18 reverse-phase HPLC using a gradient from 5 to 95% of MeCN in water supplemented with 0.1%TFA. Fractions containing pure peptides were combined and lyophilized. Peptides were stored at -20 °C until use. NMR characterization of peptides **1** and **2** is included in Supplementary Figure S1-S2. Tables S1-S2 include detailed $^{1}H$ and $^{13}C$ chemical shifts (ppm) information.

**Acyl donor glycopeptide (1)**. HR-MS: [M+H$^{+}$] Calculated for C43H76N11O17 1018.5342; found: 1018.5436.

**Table S1.** $^{1}H$ and $^{13}C$ chemical shifts (ppm) of acyl donor glycopeptide **1** at 298K.

| | GlcNAc | AA | Gly | Asn | Ala | Thr | Lys | LeuOCam | Leu |
|---|---|---|---|---|---|---|---|---|---|
| H1/C1 | 4.97/78.4 | Hα | 3.84/42.8 | 4.65/50.0 | 4.27/50.2 | 4.18/59.5 | 4.27/52.1 | 4.46/51.2 | 4.27/53.4 |
| H2/C2 | 3.73/54.4 | Hβ | | 2.71,2.76 /36.5 | 1.33/16.5 | 4.08/67.0 | 1.67,1.77 /30.3 | 1.64/39.0 | 1.54,1.61 /39.9 |
| H3/C3 | 3.53/74.4 | Hγ | | | | 1.12/18.9 | 1.36/21.9 | 1.58/24.4 | 1.54/24.3 |
| H4/C4 | 3.41/69.5 | Hδ | | | | | 1.60/26.4 | 0.85/22.3 | 0.80/20.5 |
| H5/C5 | 3.41/77.6 | Hε | | | | | 2.91/39.5 | | |
| H6/C6 | 3.67,3.79 /60.6 | $CH_2$OCam | | | | | | 4.63/63.3 | |
| NHAc | 1.97/21.8 | Ac | 1.93/22.2 | | | | | | |

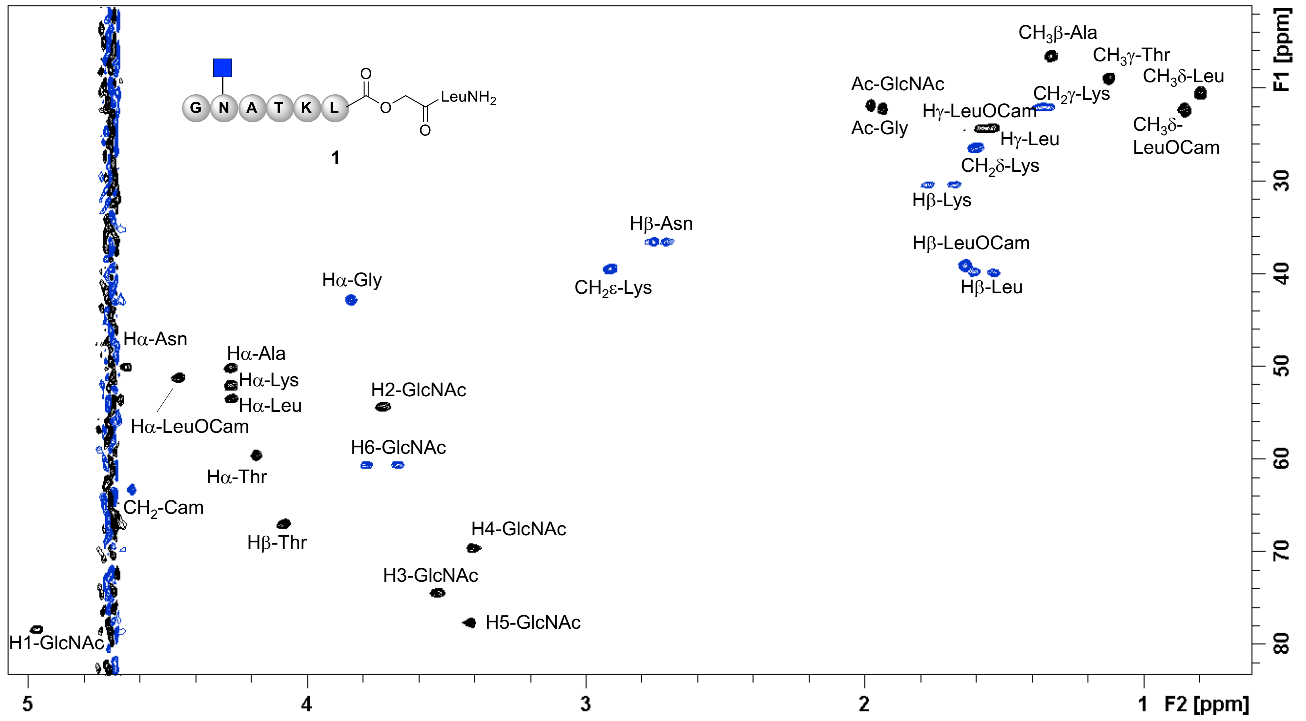


**Figure S1.** $^{13}$C-HSQC spectrum of acyl donor glycopeptide **1**. Resonance assignment has been annotated.

**Acyl acceptor peptide (2).** HR-MS: $[M+2H]^{+2}/2$ calculated for C83H139N23O27 1890.0054/2=945.0027; found: 945.0108.

**Table S2.** $^{1}$H and $^{13}$C chemical shifts (ppm) of acyl acceptor peptide **2** at 298 K.

| AA | A1 | A2/10/13 | S3/6/12/14 | D4/9 | I5 | L7/8/16 | Q11 | P15 | R17 |
|---|---|---|---|---|---|---|---|---|---|
| Hα | 3.99/49.1 | 4.51/48.6<br>4.25/50.3<br>4.22/53.1 | 4.34/55.8 | 4.47/51.8<br>4.53/51.8 | 4.09/58.9 | 4.25/52.9 | 4.18/50.5 | 4.33/60.5 | 4.22/53.1 |
| Hβ | 1.46/16.8 | 1.29/15.4<br>1.36/16.3 | 3.70/61.0 | 2.63/38.4 | 1.85/36.0 | 1.50,1.56<br>/39.5 | 1.97,2.08<br>/26.4 | 1.79,2.21<br>/29.3 | 1.62/28.1 |
| Hγ | | | | 1.13/18.8 | 0.85/14.9 | 1.55/24.2 | 2.32/31.3 | 1.93/24.7 | 1.46/24.4 |
| Hδ | | | | | 0.80/10.4 | 0.78/20.7<br>0.85/22.1 | | 3.54,3.70<br>/47.8 | 3.06/40.5 |
| Hε | | | | | | | | 2.91/47.7 | |

| AA | V18 | AA | Y19 |
|---|---|---|---|
| Hα | 3.98/59.4 | Hα | 4.46/55.0 |
| Hβ | 1.89/30.4 | Hβ | 2.81,2.98/3<br>6.3 |
| Hγ | 0.77/17.6<br>0.75/18.3 | 3,5H | 6.73/<br>115.5 |
| Hδ | | 2,6H | 7.07/<br>130.6 |
| Hε | | | |

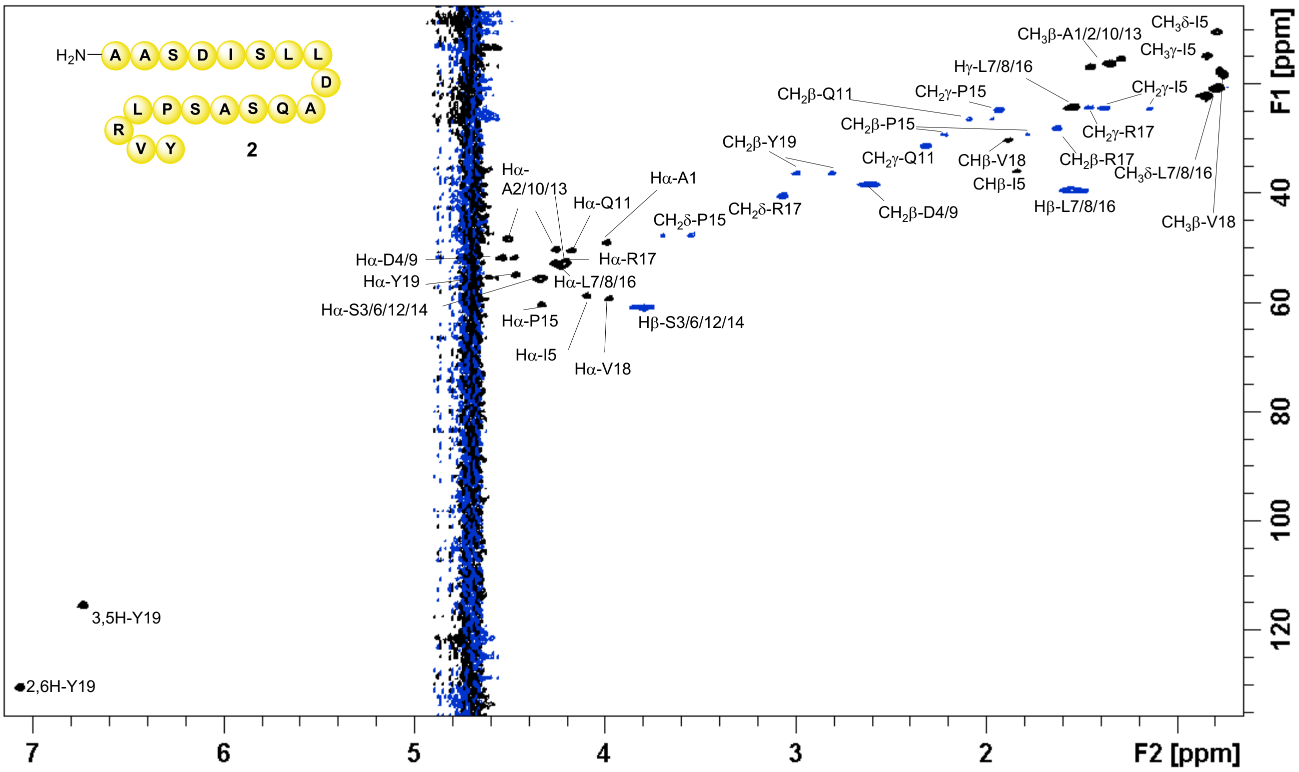


**Figure S2.** $^{13}$C-HSQC spectrum of acyl acceptor peptide **2**. Resonance assignment has been annotated.

#### 2.1.4. Procedure for glycopeptide-peptide ligation catalyzed by omniligase-1

Acyl donor glycopeptide **1** (1.6 mg, 1.6 µmol, 1 eq) was dissolved in PBS (0.64 mL, 200 mM, pH 8.0) containing DTT (10 mM). Acyl acceptor peptide **2** (4.5 mg, 2.4 µmol, 1.5 eq) was added and the resulting solution was homogenized. Omniligase-1 (0.5 nmol) was added, and the reaction mixture was incubated at RT with gently shaking. Then, 500 µL of MeCN:$H_2O$ 1:1 containing 0.1% formic acid was added and the resulting solution was lyophilized. The desired ligation product was purified by C18 reverse-phase HPLC using a gradient from 5 to 75% of MeCN in water supplemented with 0.1%TFA (60 min). Fractions containing the ligated peptide **3** were combined and lyophilized.

**Glycopeptide (3)**. HR-MS: $[M+2H]^{+}/2$ calculated for C118H198N32O41 1359.7118; found: 1359.7185.

**Table S3.** $^{1}H$ and $^{13}C$ chemical shifts (ppm) of glycopeptide **3** at 298K.

| AA | G1 | N2 | A3 | T4 | K5 | L6/13/14/22 | A7/8/16/19 | S9/12/18/20 | D10/15 |
|---|---|---|---|---|---|---|---|---|---|
| Hα | 3.84/42.7 | 4.65/49.9 | 4.28/50.0 | 4.17/59.5 | 4.25/52.3 | 4.27/52.6 | 4.51/48.4<br>4.28/50.0 | 4.34/55.7 | 4.58,4.50<br>/51.7 |
| Hβ | | 2.71,2.76<br>/36.4 | 1.34/16.4 | 4.09/66.9 | 1.67,1.75<br>/30.2 | 1.55/39.5 | 1.34/16.4<br>1.30/15.3 | 3.80/60.9 | 2.62,2.67<br>/38.1 |
| Hγ | | | | 1.13/18.8 | 1.33/21.9 | 1.56/24.3 | | | |
| Hδ | | | | | 1.63/28.0 | 0.79/20.7<br>0.85/22.2 | | | |
| Hε | | | | | 2.91/39.2 | | | | |
| Ac | 1.93/22.1 | | | | | | | | |

| AA | I11 | Q17 | P21 | R23 | V24 | AA | Y25 | GlcNAc | |
|---|---|---|---|---|---|---|---|---|---|
| Hα | 4.08/59.1 | 4.22/49.9 | 4.34/60.5 | 4.22/52.9 | 3.98/59.2 | Hα | 4.47/54.7 | H1/C1 | 4.98/78.2 |
| Hβ | 1.85/35.9 | 2.09/26.4 | 1.79,2.22<br>/29.2 | 1.63/28.0 | 1.89/30.2 | Hβ | 2.80,3.00<br>/36.4 | H2/C2 | 3.73/54.2 |
| Hγ | 1.14,1.38<br>/24.5<br>0.85/14.8 | 2.32/31.3 | 1.93/24.8 | 1.46/24.2 | 0.77/17.6<br>0.76/18.2 | 3,5H | 6.74/<br>115.4 | H3/C3 | 3.53/74.2 |
| Hδ | 0.79/10.5 | | 3.55,3.71<br>/26.4 | 3.07/40.4 | | 2,6H | 7.06/<br>130.5 | H4/C4 | 3.40/69.5 |
| Hε | | | 2.91/47.7 | | | | | H5/C5 | 3.40/77.6 |
| | | | | | | | | H6/C6 | 3.67,3.78<br>/60.5 |
| | | | | | | | | NHAc | 1.98/21.6 |

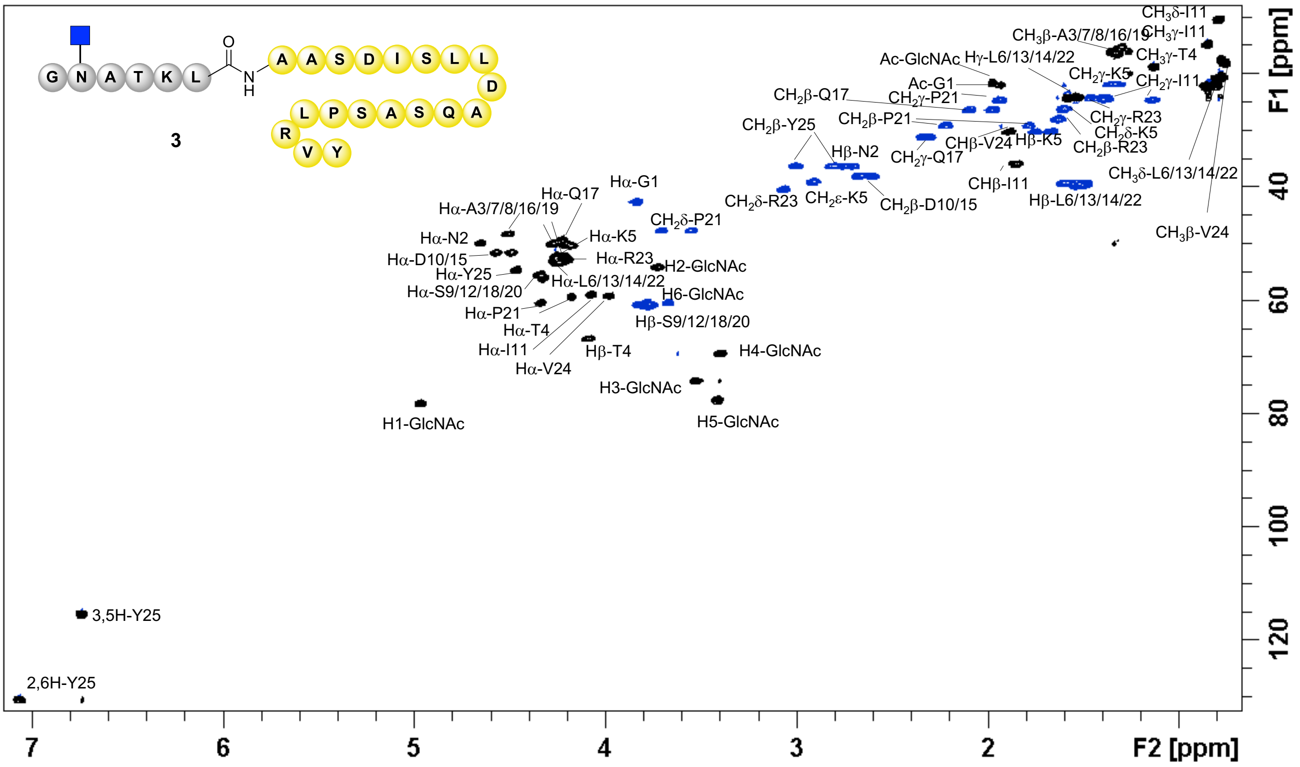


**Figure S3.** $^{13}$C-HSQC spectrum of glycopeptide **3**. Resonance assignment has been annotated.

#### 2.1.5. Synthesis of α2,6-disialo glycan derived from egg yolk and glycan oxazoline 8

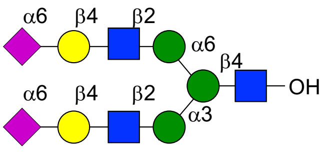


Extraction, isolation and trimming of sialylglycopeptide from egg yolk powder was performed according to our previously described procedure.[7] Briefly, commercially available egg-yolk powder (1.5 kg) was suspended in ethanol (3 L) and mechanically stirred for 15 h at RT. The filtrate was discarded, and the residue was washed twice with ethanol (500 mL). The powder was resuspended in ethanol (2 L) and stirred for 15 h. Subsequently, the suspension was filtered, and the residue washed twice with ethanol (2 x 500 mL), twice with ether (2 x 1000 mL) and dried to the air. The resulting off-white powder was suspended in aqueous ethanol (40% v/v, 2 L) and stirred for 15 h at RT. The insoluble material was discarded, and the filtrate was collected, and concentrated under reduced pressure. The combined concentrated filtrate (600 mL) was acidified to pH < 2 using trichloroacetic acid and incubate at 4 °C for 5 h. The resulting suspension was centrifuge (3000 rpm, 30 min, 4 °C) and the supernatant was filtered. The resulting clear solution was purified using an active carbon/celite column (50/50 w/w). Impurities were removed by flushing the column with water (0.1% TFA), 5% acetonitrile in water (0.1% TFA) and 15% acetonitrile in water (0.1% TFA). The glycopeptide was finally eluted with 25% acetonitrile in water (0.1% TFA). All fractions containing the product were combined, concentrated *in vacuo* to 150 mL and lyophilized. The resulting powder was purified by size exclusion chromatography (P2 biogel, Bio-Rad®) using 100 mM ammonium bicarbonate as eluent affording 1.67 g of SGP as a white powder. 100 mg of isolated SGP was dissolved in 4 mL 50 mM NaOAc pH 5.5 containing 5 mM $CaCl_2$. EndoS from *Streptococcus pyogenes* (20 µL, 2 mg/mL) and Neuraminidase from *Clostridium perfringens* (New England Biolabs #P0720L, 20 µL, 1000 units) were added and the reaction mixture was incubated for 12 h at 37 ºC. The reaction was monitored by ESI-MS and, if starting material remained another portion of EndoS and/or NeuS was added to ensure complete conversion. Enzymes were removed using an Amicon Ultra-10 centrifugal filter and the filtrate was lyophilized and purified by size-exclusion chromatography using P-2 Biogel and 100 mM ammonium bicarbonate as eluent. Fractions containing the trimmed oligosaccharide were pooled, lyophilized, and dissolved in Tris buffer (50 mM, pH 7.5) containing 10 mM $MgCl_2$. To this solution, calf intestine alkaline phosphatase (CIAP, 1% total volume, 1kU/mL), CMP-Neu5Ac (47 mg) and ST6Gal1 (1% wt/wt relative to acceptor substrate) were added, and the reaction was incubated overnight at 37 ºC. The reaction mixture was lyophilized and purified by size-exclusion chromatography using P-2 Biogel and 100 mM ammonium bicarbonate as eluent. Fractions containing α2,6-disialo glycan were pooled, and lyophilized (50 mg, 71%). $^{1}H$ and $^{13}C$ NMR analysis was consistent with previously reported data.[7] Data have been reported in Figure S4 and Table S4.

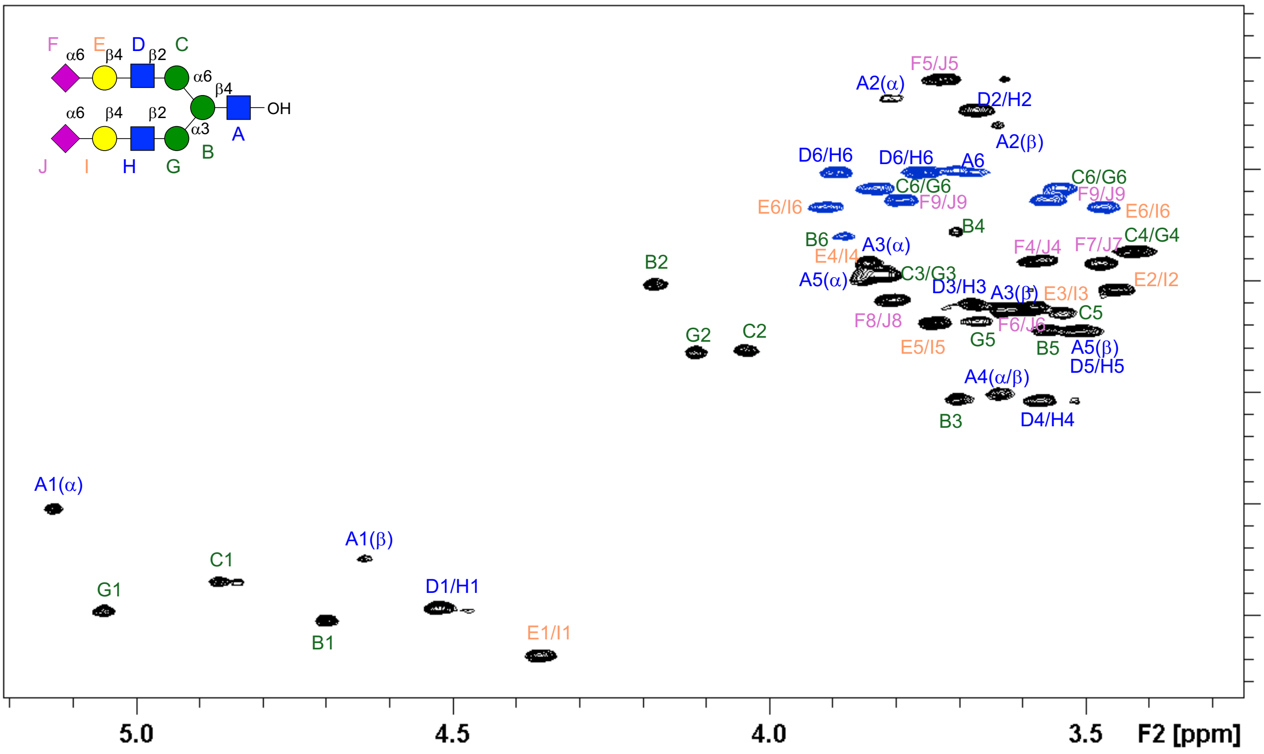


**Figure S4.** $^{13}$C-HSQC spectrum of 2,6-disialo glycan derived from egg yolk. Resonance assignment has been annotated.

**Table S4.** $^{1}$H and $^{13}$C chemical shifts (ppm) at of 2,6-disialo glycan derived from egg yolk at 298 K.

| | GlcNAcα(A) | GlcNAcβ(A) | Man(B) | Man(C) | Man(G) | GlcNAc(D/H) | Gal(E/I) | Neu5Ac(F/J) |
|---|---|---|---|---|---|---|---|---|
| H1/C1 | 5.13/90.4 | 4.64/94.9 | 4.70/100.4 | 4.87/97.0 | 5.05/99.6 | 4.52/99.3 | 4.52/99.3 | - |
| H2/C2 | 3.81/53.7 | 3.64/56.0 | 4.18/70.2 | 4.04/76.3 | 4.12/76.3 | 3.68/54.6 | 3.45/70.7 | - |
| H3/C3 | 3.84/68.3 | 3.63/72.4 | 3.70/80.5 | 3.83/69.4 | 3.83/69.4 | 3.68/72.1 | 3.58/72.4 | 2.59/39.9<br>1.65/39.9 |
| H4/C4 | 3.64/80.1 | 3.64/80.1 | 3.71/65.6 | 3.42/67.3 | 3.42/67.3 | 3.57/80.7 | 3.84/68.3 | 3.58/68.2 |
| H5/C5 | 3.85/69.7 | 3.51/74.4 | 3.56/74.4 | 3.54/72.8 | 3.67/73.5 | 3.51/74.4 | 3.74/73.7 | 3.73/51.8 |
| H6/C6 | 3.67/60.2 | 3.67/60.2 | 3.88/65.9<br>3.71/65.9 | 3.83/61.7<br>3.54/61.6 | 3.83/61.7<br>3.54/61.6 | 3.89/60.2<br>3.76/60.2 | 3.91/63.4<br>3.47/63.3 | 3.63/72.4 |
| H7/C7 | - | - | - | - | - | - | - | 3.47/68.3 |
| H8/C8 | - | - | - | - | - | - | - | 3.80/71.7 |
| H9/C9 | - | - | - | - | - | - | - | 3.79/62.8<br>3.56/62.7 |
| NHAc | 1.98/22.3 | 1.98/22.3 | - | - | - | 1.95/22.0 | - | 1.98/22.3 |

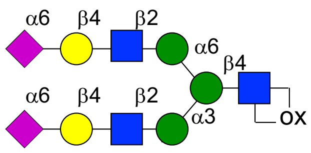


α2,6-disialo biantennary *N*-glycan (100 mg, 0.05 mmol) was dissolved in water (5 mL) and triethylamine (140 µL, 1.0 mmol) was added. The reaction mixture was suspended in an ice bath and stirred at 0 °C for 5 min. 2-Chloro-1,3-dimethylimidazolinium chloride (35 mg, 0.5 mmol) was added and the reaction mixture was stirred at 0 °C for 2 h. Complete conversion into oxazoline was assessed by ESI-MS. The reaction mixture was lyophilized and purified by size exclusion chromatography using P2 gel (column height 15 cm). Subsequent lyophilization afforded oxazoline **8** as a white solid (70 mg, 71%) that was stored as 10 mM NaOH solution at -80 ºC. $^1$H NMR analysis was consistent with previously reported data.[2] 1H NMR (400 MHz, d2o) δ 5.95 (d, J = 7.3 Hz, 1H), 4.98 (s, 2H), 4.82 (s, 2H), 4.46 (d, J = 8.2 Hz, 4H), 4.32 – 4.22 (m, 5H), 4.03 (d, J = 14.2 Hz, 7H), 3.88 – 3.24 (m), 2.84 (d, J = 2.2 Hz, 2H), 1.96 – 1.83 (m, 24H), 1.57 (t, J = 12.1 Hz, 4H).

### 2.2. Protein Expression and Purification

#### 2.2.1. Expression and purification of recombinant human Galectin-3 (CRD)

The gene encoding the carbohydrate recognition domain (CRD) of human Galectin-3 (*h*Gal3-CRD) (amino acid residues 114-248) was inserted into the pET21a expression vector. The plasmid was synthesized by Genscript. Transformation of BL21 (D3) *E. coli* competent cells with the expression vector was performed by heat shock method (42 °C for 90 sec, 5 min in ice). After being incubated overnight on agar plates in the presence of ampicillin at 37 °C, one single colony harbouring the expression construct was inoculated into 5 mL Luria Broth (LB) medium containing 100 µg/mL ampicillin and it was cultured overnight at 37 °C with shaking. A certain volume of the pre-culture was added to 2 L of new LB medium with ampicillin. Cells were grown at 37 ºC until an OD600 of 0.6-1.2. Then, protein expression was induced with 1 mM Isopropyl β-D-1-thio-galactopyranoside (IPTG) and cell culture grow for another 3 h at 37 ºC. Afterwards, cells were harvested by centrifugation at 5500 rpm for 20 min. The pellet was suspended in 10 mL lysis buffer (22 mM Tris-HCl pH 7.5, 5 mM EDTA and 1 mM DTT) and the cell suspension was lysed by sonication (8 x 30 sec, with 1 min intervals between each burst). The crude extract was clarified by centrifugation at 35000 rpm for 30 min. The soluble fraction was loaded onto 5 mL α-Lactose-Agarose resin (Sigma-Aldrich) equilibrated with PBS pH 7.8. The column was washed with 50 mL of buffer equilibration. The recombinant hGal3-CRD was eluted with approximately 7 mL (150 mM α-Lactose pH 7.4 in PBS) and the protein purity was checked by 4-12% SDS-PAGE as one clear band, the position of which corresponds well to the calculated molecular weight of 15 kDa. The identity of the protein was further confirmed by mass spectrometry.

#### 2.2.2. Expression and purification of recombinant human Interleukin-18 (6)

The DNA encoding the sequence of human interleukin-18 (IL-18) (UniprotKB Q14116, residues 37–193) carrying a C-terminal Strep tag after a HRV3C cleavage site and a N-terminal 8xHis tag before a Tobacco Etch Virus (TEV) cleavage site was subcloned between AvaI and EcoRI restriction sites into pMAL-c4x-H vector (Genscript) and codon optimized for expression in *E. coli.* The plasmid was synthesized by Genscript. Transformation of BL21 (D3) *E. coli* competent cells with the expression vector was performed by heat shock method (42 °C for 10 sec, 5 min in ice). After being incubated overnight on agar plates in the presence of ampicillin at 37 °C, one single colony harbouring the expression construct was inoculated into 5 mL 2xYT medium containing 100 µg/mL ampicillin and it was cultured overnight at 37 °C with shaking. 4 mL of the pre-culture was added to 2 L of new 2x YT medium with ampicillin. Cells were grown at 37 ºC until an OD600 of 0.6-0.8 was reached. Then, protein expression was induced with 1 mM IPTG and cell culture grow overnight at RT. Afterwards, cells were harvested by centrifugation at 5500 rpm for 20 min at 4 ºC. The pellet was suspended in 100 mL lysis buffer (50 mM Tris-HCl pH 8.0, 250 mM NaCl, 10 mM imidazole, Triton-X 0.1%, lysozyme 1 mg/mL) and cells were lysed by sonication (3x10 sec, with 10 sec intervals between each burst). The crude extract was clarified by centrifugation at 35.000 rpm for 20 min at 4 ºC. The soluble fraction was loaded onto 5 mL Ni-NTA Sepharose column equilibrated with buffer A (50 mM Tris-HCl pH 8.0, 250 mM NaCl, 10 mM imidazole). The column was washed with 50 mL of buffer A, followed by 50 mL of buffer B (50 mM Tris-HCl pH 8.0, 250 mM NaCl, 50 mM imidazole). The recombinant protein was eluted with approximately 15 mL of elution buffer (50 mM Tris-HCl pH 8.0, 250 mM NaCl, 250 mM imidazole) and the protein purity was checked by 4-12% SDS-PAGE as one clear band, the position of which corresponds well to the calculated molecular weight of 67.7 kDa. The sample was loaded onto 5 mL amylose resin equilibrated with 20 mM Tris-HCl, 200 mM NaCl, 1 mM EDTA, pH 7.4 (Column Buffer). The column was washed with 50 mL of Column Buffer and the fusion protein was eluted with 15 mL of Column Buffer containing 10 mM maltose. The protein-containing fractions were pooled, and cleavage of the fusion protein was performed by incubation with TEV protease (one unit per 2 µg of fusion protein) at 4 ºC overnight. Finally, the digest was loaded onto Strep-

Tactin resin (2 mL) equilibrated with washing buffer (100 mM Tris-HCl pH 8.0, 150 mM NaCl, 1 mM EDTA). After washing the column with 50 mL of the same buffer the target protein was eluted with 15 mL of washing buffer containing 2.5 mM desthiobiotin. The protein purity was checked by 4-12% SDS-PAGE as one clear band, the position of which corresponds well to the calculated molecular weight of 23 kDa. The identity of the protein was further confirmed by mass spectrometry.

#### 2.2.3. Expression and purification of recombinant human Interferon alpha-2a (7)

The DNA encoding the sequence of human interferon alpha-2a (IFNα-2a) (UniprotKB P01563, residues 24–188) including an additional Phe residue was subcloned as an N-terminal His-tagged SUMO fusion protein (also including a TEV cleavage site) into the pET11a vector (Genscript) and codon optimized for expression in *E. coli*. The plasmid was synthesized by Genscript. Transformation of BL21 (D3) *E. coli* competent cells with the expression vector was performed by heat shock method (42 °C for 10 sec, 5 min in ice). After being incubated overnight on agar plates in the presence of ampicillin at 37 °C, one single colony harbouring the expression construct was inoculated into 5 mL 2xYT medium containing 100 µg/mL ampicillin and it was cultured overnight at 37 °C with shaking. 2 mL of the pre-culture was added to 1 L of new 2x YT medium with ampicillin. Cells were grown at 37 ºC until an OD600 between 0.6-0.8 was reached. Then, protein expression was induced with 1 mM IPTG and the cell culture was grown overnight at RT. Afterwards, cells were harvested by centrifugation at 5500 rpm for 20 min at 4 ºC. The pellet was suspended in 50 mL lysis buffer (50 mM Tris-HCl pH 8.0, 250 mM NaCl, 10 mM imidazole, Triton-X 0.1%, lysozyme 1mg/mL) and cells were lysed by sonication (3x10 sec, with 10 sec intervals between each burst). The crude extract was clarified by centrifugation at 35000 rpm for 20 min at 4 ºC. The soluble fraction was loaded onto 5 mL Ni-NTA Sepharose column equilibrated with buffer A (50 mM Tris-HCl pH 8.0, 250 mM NaCl, 10 mM imidazole). The column was washed with 50 mL of buffer A, followed by 50 mL of buffer B (50 mM Tris-HCl pH 8.0, 250 mM NaCl, 50 mM imidazole). The recombinant protein was eluted with approximately 15 mL of elution buffer (50 mM Tris-HCl pH 8.0, 250 mM NaCl, 250 mM imidazole). Fractions containing the protein were joined, concentrated, dialyzed against 50 mM Tris-HCl pH 8.0, 250 mM NaCl, and submitted to digestion with TEV protease (one unit per 2 µg of fusion protein) at 4 ºC overnight. The cleaved $His_8$-SUMO tag and TEV protease were separated from the pure target protein using Ni-NTA affinity chromatography. Briefly, the digest was loaded onto Ni-NTA Sepharose column equilibrated with buffer A and washed with an additional 20 mL. Untagged IFNα-2a was collected in the flow-through and wash steps. The protein purity was checked by 4-12% SDS-PAGE as one clear band, the position of which corresponds well to the calculated molecular weight of 19.4 kDa. The identity of the protein was further confirmed by mass spectrometry.

### 2.3. Chemoenzymatic Protein Modification

#### 2.3.1. General methods

*a MALDI-TOF protein analysis*

All preparations were desalted using ZipTip® C4 micro-columns (Millipore) (2 µL samples) and eluted using 0.5 µL SA matrix (Sinapinic acid, 10 mg/mL in 70:30 MeCN:MQ supplemented with 0.1% TFA) or HCCA matrix (α-cyano-4-hydroxycinnamic acid, 10 mg/mL in 70:30 MeCN:MQ supplemented with 0.1% TFA) onto a GroundSteel massive 384 target (Bruker Daltonics).

Mass spectrometry: an AutofleXtreme MALDI-TOF/TOF spectrometer (Bruker Daltonics) was used in linear mode with the following settings: 5000-40000 Th window, linear positive mode, ion source 1: 20 kV, ion source 2: 18.5 kV, lens: 9 kV, pulsed ion extraction of 120 ns, high gating ion suppression up to 1000 Mr. Mass calibration was performed externally with protein 1 standard calibration mixture (Bruker Daltonics) in the same range as the samples. Data acquisition was performed using FlexControl 3.4 software (Bruker Daltonics), and peak peaking and subsequent spectra analysis was performed using FlexAnalysis 3.4 software (Bruker Daltonics).

*b UHPLC-HRMS protein analysis*

Protein purity and identity was assessed by intact protein mass spectrometry. All samples were desalted. To this aim, 2 µL of protein samples were analyzed on an Agilent 6560 Ion Mobility LC/Q-TOF mass spectrometer. A 15 min LC gradient from 20% to 80% buffer B (Buffer A: MQ supplemented with 0.1% FA; buffer B: 70% isopropanol, 20% MeCN, 10% MQ supplemented with 0.1% FA) at a flow rate of 0.3 mL/min was used to separate protein species. A Poroshell 120 EC C18 (50 mm × 2.1 mm, 2.7 µm, Agilent Technologies) analytical column, thermostated at 50 ºC, was used for protein separation. Deconvoluted spectra were obtained using Bruker Daltonics Maximum Entropy deconvolution software with a mass range of 5 to 30 kDa, data point spacing of 0.1 m/z and maximum resolving power at high resolution as settings.

#### 2.3.2. Activity assays of peptiligases in glycopeptide-protein ligations - general procedure

Purified protein was diluted to 150 µM in 100 mM tricine, pH 8.0 containing 2 equivalents of the peptide to be conjugated. The appropriate peptiligase (2 µM) was added and the reaction mixture was incubated at RT with gently shaking for 2 h. 10 µL aliquote was treated with MQ containing 0.1% formic acid, and reaction conversion was estimated by UHPLC-HRMS. Conversions obtained for BSA and *h*Galectin-3 are included in Table S5 as derived from the corresponding deconvoluted mass spectra (Figure S5).

**Table S5.** Evaluation of peptidoligase-mediated protein-glycopeptide ligations.

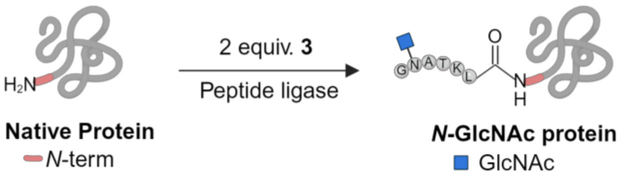


| Native Protein (N-term AA) | Equiv. 3 | Peptiligase | Conversion (%)[a] |
|---|---|---|---|
| BSA (DT) | 2 | Omniligase-1 | 8 |
| BSA (DT) | 2 | Thymoligase | 35 |
| *h*Gal-3 (ML) | 2 | Omniligase-1 | 20 |
| *h*Gal-3 (ML) | 20 | Thymoligase | 51 |

[a] Conversions were estimated by mass spectrometry analysis and referred to the limiting reagent.

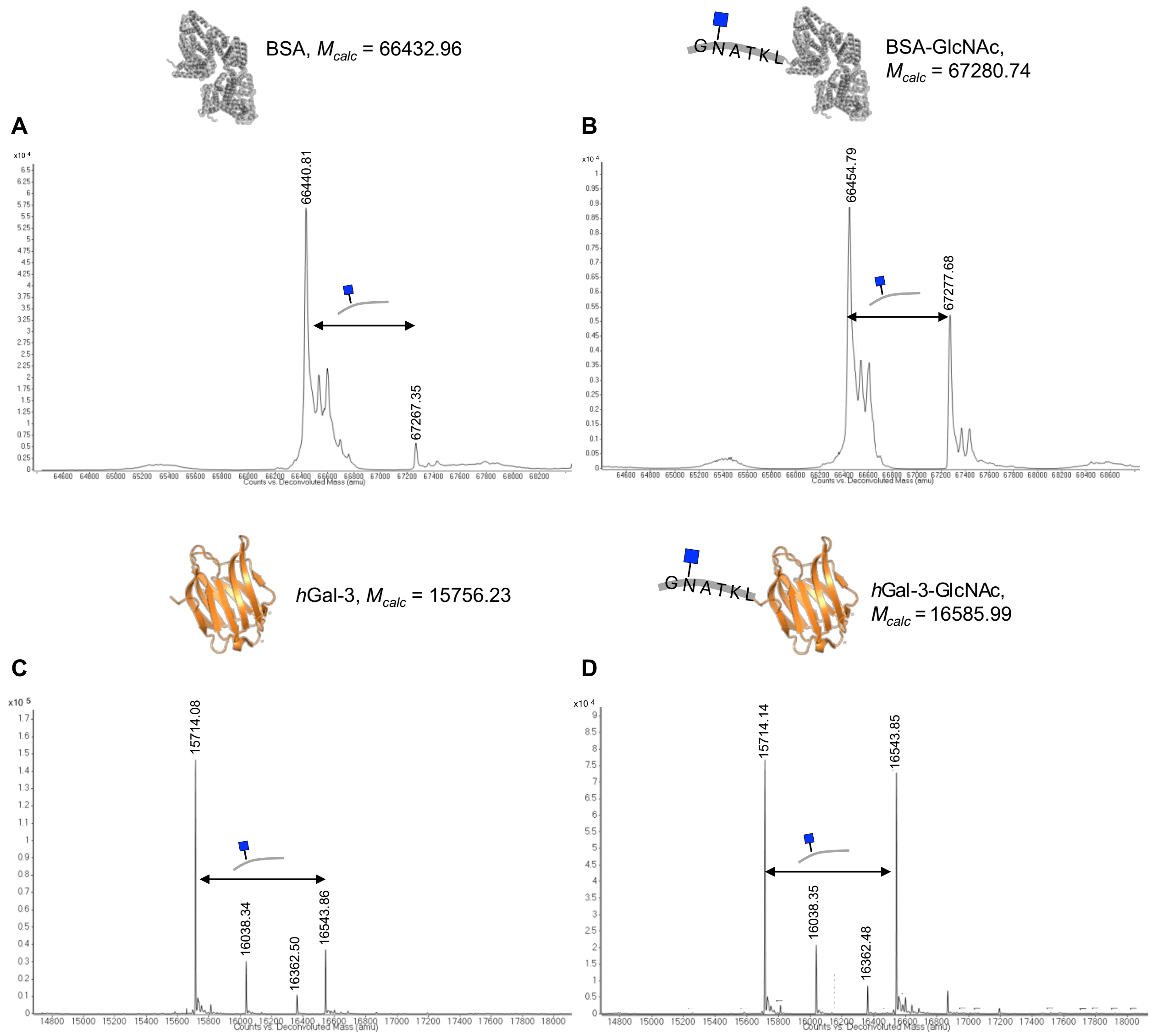


**Figure S5.** Deconvoluted mass spectra of peptiligase-catalyzed crude reactions of glycopeptide **3** ligation. A) Omniligase-1 and B) Thymoligase were tested as enzymes for BSA glycotagging. Similar reactions in the presence of C) Omniligase-1 and D) Thymoligase were performed for *h*Gal-3.

##### 2.3.3. Glycopeptide-protein ligation. Procedure for the preparation of glycotagged proteins IL-18-GlcNAc (6a) and IFNa-2a (7a)

Purified protein was diluted to 150 µM in 100 mM tricine, pH 8.0 containing 5 equivalents of the peptide to be conjugated. Thymoligase (1 mol%) was added and the reaction mixture was incubated at RT with gently shaking. Reaction progress was monitored by QTOF-MS and if unmodified protein remained after 2 h, a second portion of glycopeptide **3** was added and allowed to react for additional 2 h. Next, thymoligase was removed by affinity chromatography techniques. Briefly, reaction samples were loaded into a Ni-NTA Sepharose column equilibrated with buffer A (50 mM Tris-HCl pH 8.0, 150 mM NaCl). The column was washed with 10 column volumes of buffer A. IL-18 and IFNα-2a derived proteins were collected in the flow-through and washing steps, while thymoligase was retained in the resin. In addition, IL-18 modified protein, was further purified by Strep-tag affinity chromatography (as described in Section 2.2.2). Fractions containing the desired proteins were combined and concentrated using a 10 kDa centrifugal filter. The identity and purity of the resulting proteins was checked by 4-12% SDS-PAGE and ESI-QTOF-MS (Figures S6 and S7).

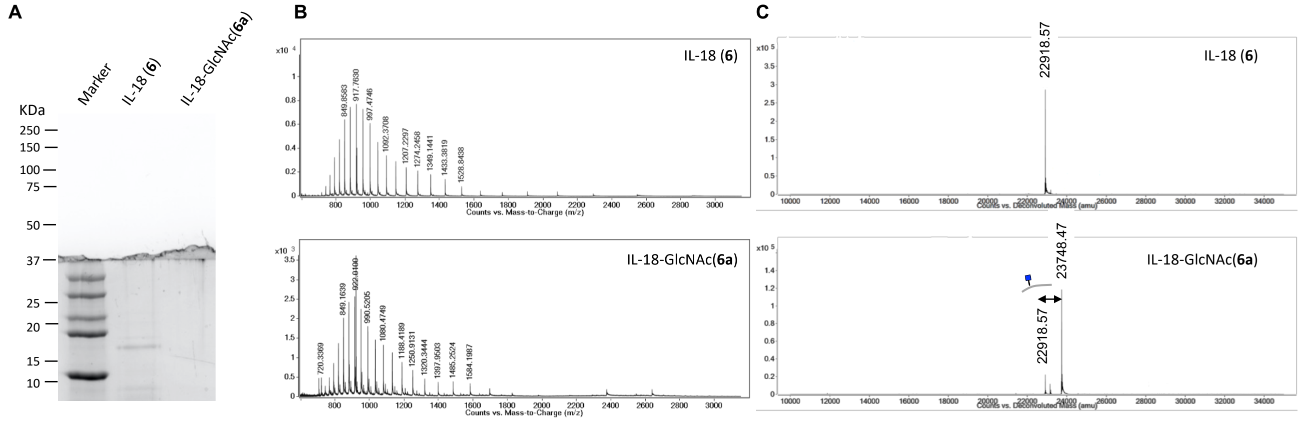


**Figure S6.** A) SDS-PAGE (4-12%, 200V, 40min, stained with Coomassie) of IL-18 (**6**) and glycovariant IL-18-GlcNAc (**6a**). B) ESI-QTOF spectra of **6** and **6a**. C) Deconvoluted ESI-QTOF spectra of **6** and **6a**.

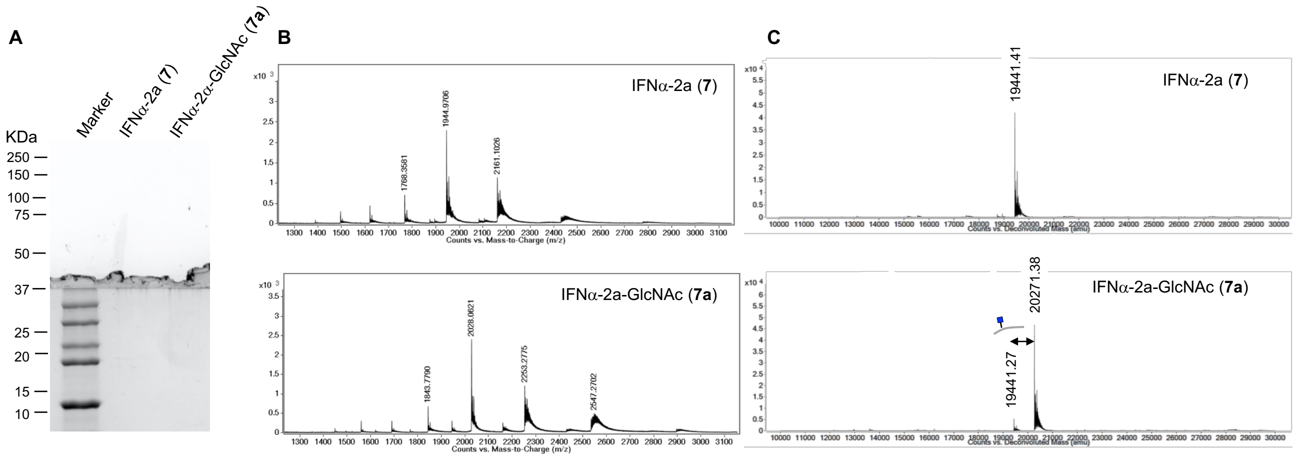


**Figure S7.** A) SDS-PAGE (4-12%, 200V, 40 min, stained with Coomassie) of IFNα-2a (**7**) and glycovariant IFNα-2a -GlcNAc (**7a**). B) ESI-QTOF spectra of **7** and **7a**. C) Deconvoluted ESI-QTOF spectra of **7** and **7a**.

#### 2.3.4. Glycopeptide-protein ligation. Procedure for the preparation of insulin glycolvariants 9a and 9b

Human insulin was diluted to 150 μM in 100 mM tricine, pH 8.0 containing 2 equiv. of the peptide **1** to be conjugated. The appropriate peptiligase (1.5 mol%) was added and the reaction mixture was incubated at RT with gently shaking. Reaction progress was monitored by MALDI-TOF MS and if unmodified protein remained after 2 h a second portion of glycopeptide **3** was added and allowed to react for additional 2 h. Then, MQ containing 0.1% formic acid was added and the resulting solution was lyophilized. The desired protein bioconjugation product was purified by C18 reverse-phase HPLC using a gradient from 5 to 50% of MeCN in water supplemented with 0.1% TFA (110 min). Fractions containing modified protein were combined and lyophilized. Modified proteins were stored at -20 °C until further use. Analysis of the sample by ESI-QTOF-MS revealed product purity of > 95% (Figure S8).

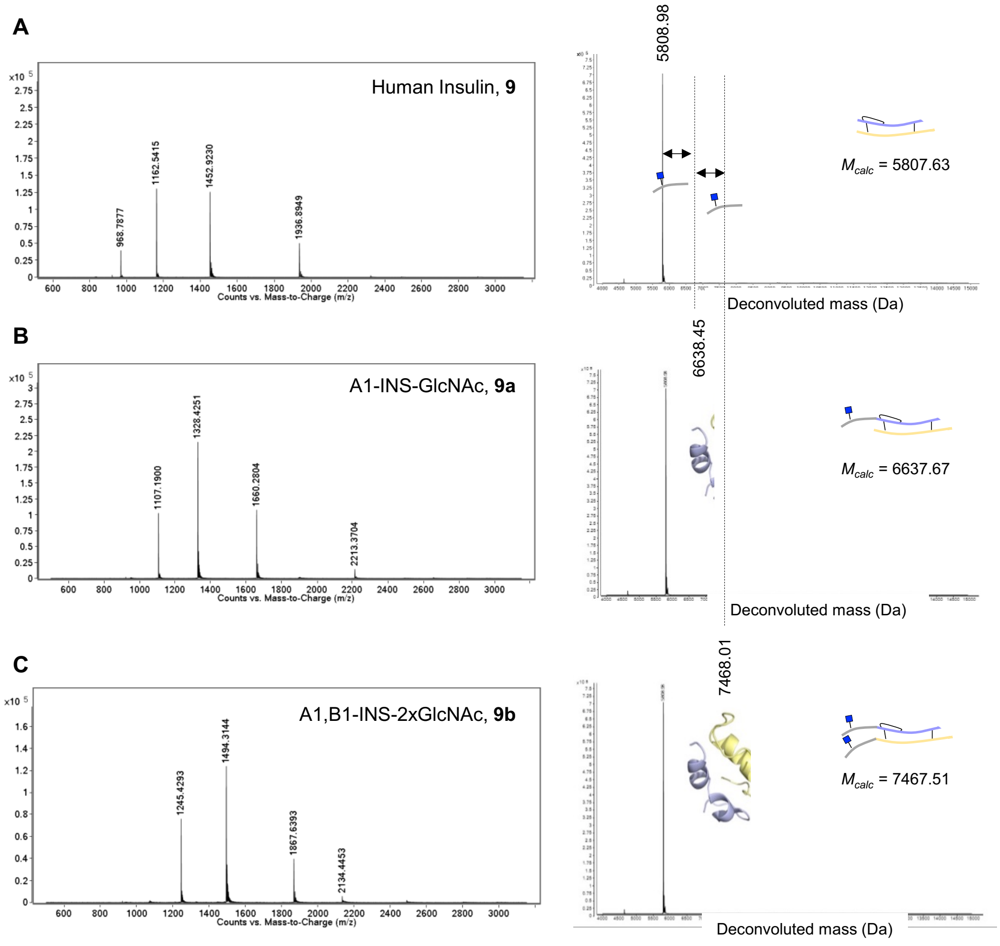


**Figure S8.** ESI-QTOF and deconvoluted ESI-QTOF spectra of a) recombinant human insulin **9** (as standard), b) Ac-GN(GlcNAc)ATKL-INS **9a** and c) 2x(Ac-GN(GlcNAc)ATKL)-INS **9b**.

*Site-Selectivity of Omniligase-1-catalyzed Peptide Ligation of Human Insulin*

Human insulin consists of two peptide chains, A (amino acids A1–A21) and B (amino acids B1–B30), which are connected by two disulfide bonds (A7–B7 and A20–B19). Therefore, insulin contains two potential ligation sites, chain A and B *N*-terminus. Chain A and B have Gly-Ile and Phe-Val at the N-terminus which based on PILS (proteomic identification of ligation sites)[8] are expected to be appropriate substrates of peptiligases. To determine the site-selectivity of omniligase-1-catalyzed ligation using glycopeptide **3**, analysis of the peptides derived from the monoligated insulin **9a** was carried out using mass spectrometry techniques. Thus, a solution of **9a** in TRIS 20 mM pH 8 containing 5 mM DTT was incubated at RT. After 30 min the reaction mixture was analyzed by MALDI-TOF spectrometry using HCCA (α-cyano-4-hydroxycinnamic acid) as matrix.

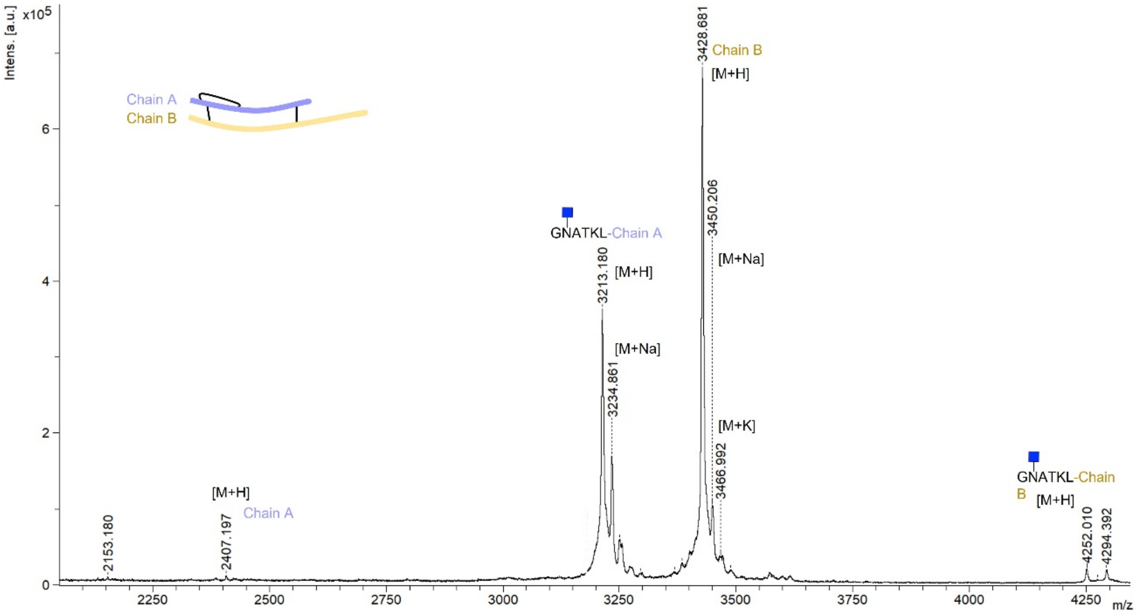


**Figure S9.** MALDI-TOF spectrum of peptides derived from mono-ligated insulin **9a** after treatment with DTT. Glucosylated chain A and unmodified chain B were detected as the major species showing the selectivity of the reaction towards ligation of chain A.

#### 2.3.5. Transglycosylation reactions mediated by EndoCC1-N180H. Installation of disialylated complex *N*-glycan

*a. Preparation of IL-18-S2G2(**6b**) and IFNα–2a-S2G2 (7**b**) variants*

A mixture of of glucosylated protein (**6a**/**7a**, 30-50 μM), S2G2 oxazoline **8** (40 equiv.), and the N180H mutant of EndoCC1 (0.5 mol%) was incubated in Tris-HCl buffer (20 mM, pH 7.5) at RT. Consumption of oxazoline **8** was monitored by LC-MS analysis while production of disialylated proteins (**6b**/**7b**) was ascertained by ESI-QTOF-MS. After 1 h, an additional portion of oxazoline **9** was added to drive the reaction towards completion. Subsequently, the EndoCC1-N180H and the protein variants were separated by different chromatography techniques. Briefly, IL-18-S2G2 (**6b**), carrying a C-terminal Strep-tag was purified by Strep-tag affinity chromatography (as described in Section 2.2.2). Fractions containing the desired protein were combined and concentrated using a 10 kDa centrifugal filter. The identity and purity of the resulting IL-18 derived protein was checked by 4-12% SDS-PAGE and ESI-QTOF-MS analysis (Figure S10 and S11).

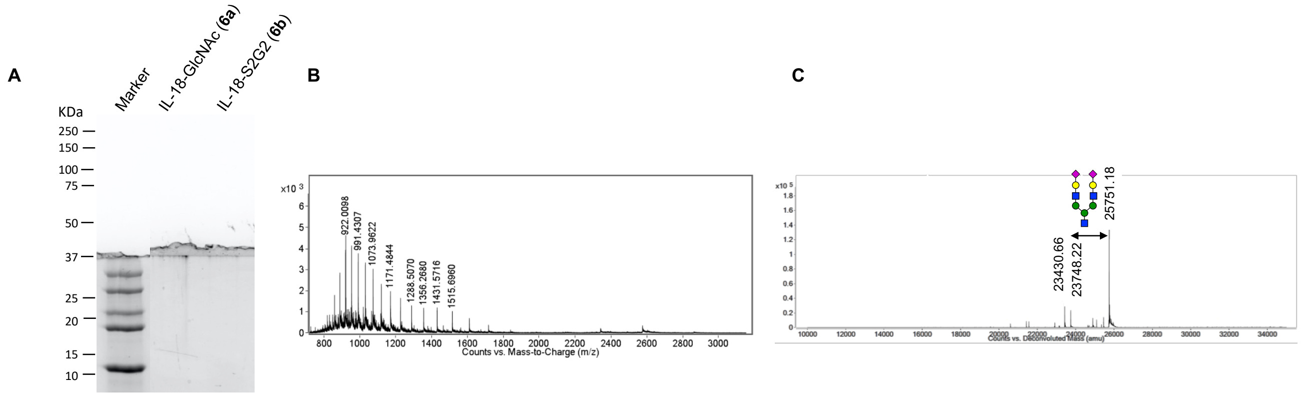


**Figure S10.** A) SDS-PAGE (4-12%, 200 V, 40 min, stained with Coomassie) of IL-18-GlcNAc (**6a**) and glycovariant IL-18-S2G2 (**6b**). B) ESI-QTOF and C) Deconvoluted ESI-QTOF spectra of **6b**.

An alternative purification strategy was implemented for IFNα-2a-S2G2 (**7b**) since no-affinity tag was present on the protein construct. Briefly, EndoCC1-N180H, carrying a His8-tag, was removed by Ni-NTA affinity chromatography. The collected flow-through, containing the protein of interest was loaded onto 2 mL Concanavalin A (Con A) agarose resin equilibrated with 1 M NaCl, 5 mM $MgCl_2$, 5 mM $MnCl_2$, 5 mM $CaCl_2$, pH 7.5 (equilibration buffer) and then with 20 mM Tris, 500 mM NaCl pH7.4 (binding buffer). The column was washed with 10 mL of binding buffer and the protein of interest was eluted with binding buffer supplemented with 500 mM D-glucose. Fractions containing protein were combined and concentrated using a 10 kDa centrifugal filter. Protein purity was checked by 4-12% SDS-PAGE and ESI-QTOF-MS and showed IFNα-2a-S2G2 (**7b**) as the main species (above 75% quantified by mass spectrometry analysis) (Figure S11).

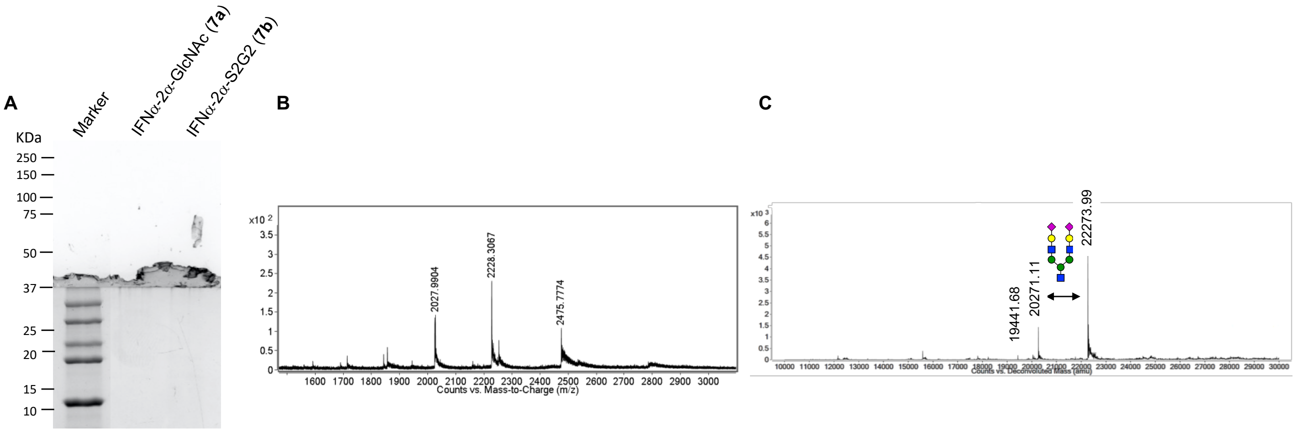

**Figure S11.** A) SDS-PAGE (4-12%, 200 V, 40 min, stained with Coomassie) of IFNα-2a-GlcNAc (**7a**) and glycovariant IFNα-2a-S2G2 (**7b**). B) ESI-QTOF and C) Deconvoluted ESI-QTOF spectra of **7b**.

Further purification of the final protein construct (IFNα-2a-S2G2, **7b**) was carried out by size exclusion chromatography. Briefly, a Reprosil 125 SEC column (MW range: 5-100 kDa, particle size 3 μm, 300 mm length and 4.6 mm inner diameter) was equilibrated with 50 mM sodium phosphate, 150 mM NaCl, pH 7.0. 50 μL sample at 0.23 mg/mL was loaded, and the protein was purified using an isocratic elution at 0.3 mL/min (see Figure S12).

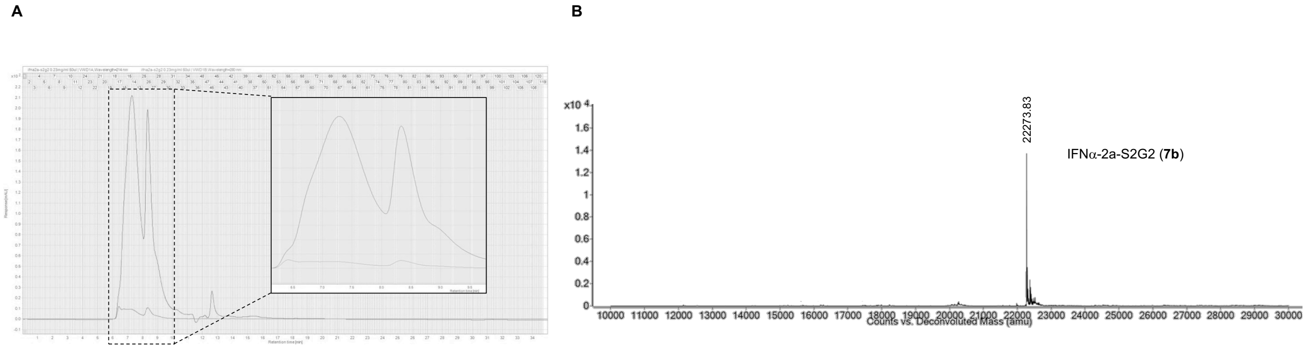

**Figure S12.** A) HPLC profile of IFNα-2a-S2G2, **7b** using Reprosil 125 SEC chromatography. Two peaks were detected. While no IFNα-2a derived protein was detected between 6.5-8 min elution time, the peak at 8.25 min corresponded to product **7b**, as indicated by mass spectrometry analysis. B) Deconvoluted mass spectrum of sample eluting at 8.25 min.

*b. Preparation of glycoinsulin variants **9c** and **9d***

A mixture of the glucosylated protein (128 µg, 1.3 mg/mL), S2G2 oxazoline 8 (20 equiv.), and the N180H mutant of EndoCC1 (2 µg, 0.018 µg/µL) was incubated in Tris-HCl buffer (20 mM, pH 7.5) at RT. The reaction was monitored by LC-MS analysis. After 2 h, an additional portion of oxazoline **8** was added to drive the reaction towards completion. Then, the sialylated insulin product was isolated by C18 reverse-phase HPLC using a gradient from 5 to 50% of MeCN in water supplemented with 0.1%TFA (110 min). The identity and purity of the resulting glycosylated insulins **9b** and **9d** was checked by ESI-QTOF-MS analysis (Figure S13).

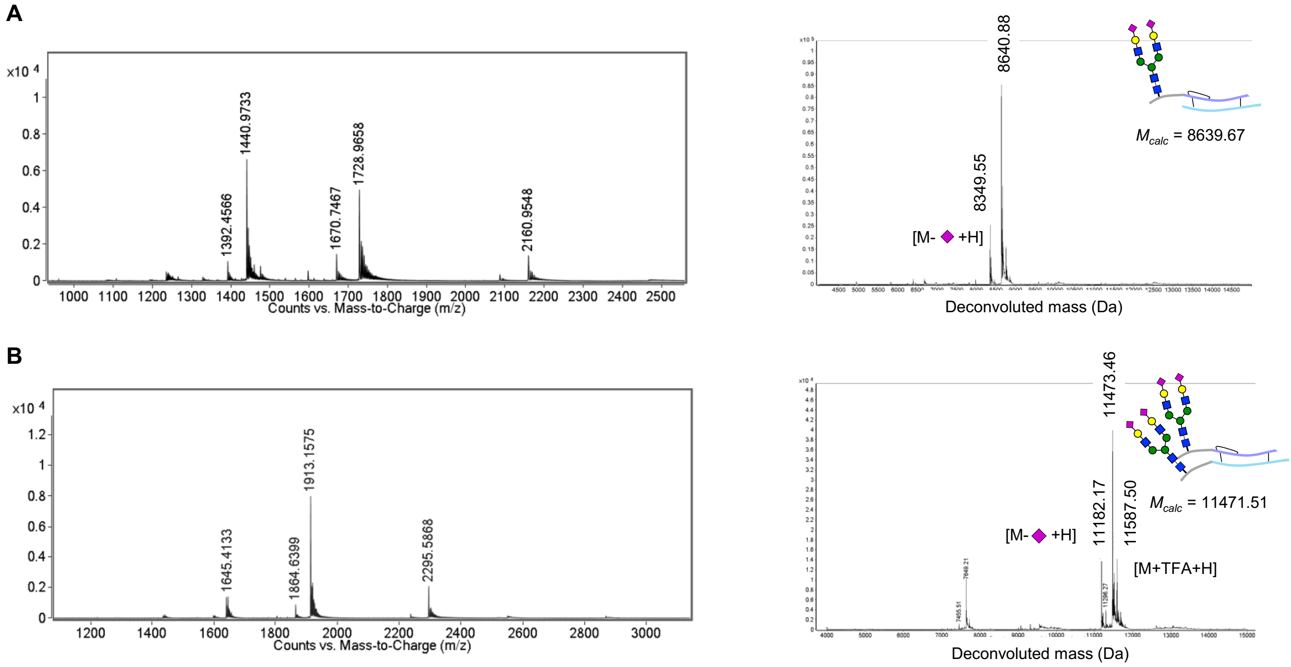


**Figure S13.** ESI-QTOF and deconvoluted ESI-QTOF spectra of A) A1-INS-S2G2 **9c** and B) A1,B1-INS-2xS2G2 **9d**.

### 2.4. Structural and Biophysical Analysis of Insulin Variants

#### 2.4.1. NMR experiments

*General remarks.* All NMR experiments were performed using a Bruker AVANCE 2 600 MHz spectrometer equipped with standard triple-channel probe. Typical concentrations of proteins were 50-100 µM and phosphate buffered saline solutions were employed.

*2D-DOSY experiments.* Samples for DOSY experiments were prepared in phosphate buffer (sodium phosphate 10 mM, NaCl 150 mM, pH 1.6, prepared using 90:10 $H_2O$:$D_2O$) with a protein concentration of 50 μM. DOSY experiments were measured at 25 °C and performed employing an array of 16 spectra for each experiment (256 or 512 transients each, with 3 sec recycle delay), and varying the gradient strength between 5% and 95%. The lengths of and delays between the gradient pulses were optimized for each sample (Figure S14). Data fitting and diffusion coefficients determination were performed employing the T1/T2 Relaxation module available in the Bruker TopSpin software. RNAse A was used as protein reference and its diffusion coefficient was calculated with model coordinates (PDB 3rn3) using the program HYDROPRO.[9]The buffer properties (viscosity) were tuned to match the computed result with the experimental value obtained for RNAse A and used in the simulations. Similarly, diffusion coefficients for glycoinsulin variants **9a-9d** were predicted based on the models generated using modelling procedures (see Section 2.4.3 for details).

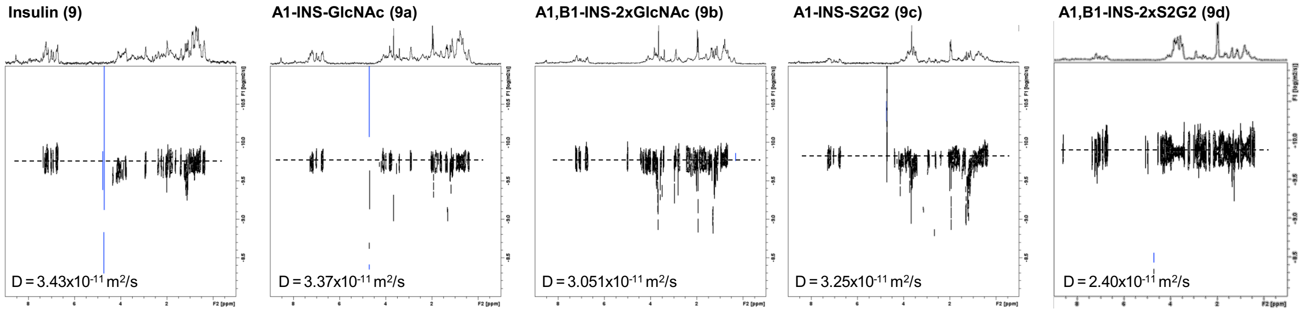


**Figure S14.** 2D-DOSY NMR spectra of human insulin **9** and glycovariants **9a-9d**.

*1H-NMR experiments.* Samples for the 1D 1H-NMR experiments were prepared in phosphate buffer (sodium phosphate 10 mM, NaCl 150 mM, pH 1.6, prepared using 90:10 $H_2O$:$D_2O$) with a protein concentration of 50 μM. Samples were prepared at RT and maintained at 60 °C. The aggregation process was monitored by following changes on the $^1H$ signal intensities of the protein. All $^1H$ spectra were acquired with 128 scans and 2 sec recycle delay. The excitation sculpting scheme was employed to suppress $H_2O$ signal.[10]

#### 2.4.3. Molecular modelling and MD simulations of glycoinsulins 9a-9d

Globular structures of modified insulin sequence (incorporating GNATKL extra sequence at the *N*-terminal end of chain A and both chains A and B) were built using AlphaFold2.[11] The top-ranked structures (predicted with the highest confidence, i.e., with the highest global pLDDT) were chosen for further elaboration. Next, GlcNAc or the disialoglycan complex *N*-glycan were incorporated into the protein structures using the glycoprotein builder module available in the GLYCAM web portal, www.glycam.org. Structures were used as initial coordinates for MD simulations. MD simulations were performed using Amber16 program with the GLYCAM06j-1 and leaprc.protein.ff14SB force field parameters. Thereafter, the starting 3D geometries were placed into a 12Å octaedral box of explicit TIP3P waters, and counterions were added to maintain electroneutrality. Na+ cations were introduced since they are used experimentally. Two consecutive minimizations were performed: 1) involving only the water molecules and ions, and 2) involving the whole system. Molecular dynamics simulations without constraints were recorded. The system was then heated and equilibrated in two steps: 1) 20 ps of MD heating the whole system from 0 to 300 K, followed by 2) equilibration of the entire system during 100 ps at 300 K. The equilibrated structures were the starting points for MD simulations (500 ns) at constant temperature (300 K) and pressure (1 atm). A detailed analysis of each MD trajectory (for example r.m.s.d. evaluation, dihedral angles and radius of gyration) was accomplished using the cpptraj module included in Amber-Tools 16 package.

#### 2.4.5. Transmission electron microscopy

Aliquots (100 µL) of 30 µM insulin samples **9** and **9a-9d** in 10 mM sodium phosphate at pH 1.6 were incubated in polypropylene Eppendorf tubes at 60 ºC for 48 h. Aliquots (10 µL) of insulin **9a-9d** solutions were adsorbed onto glow-discharged carbon-coated copper grids and stained with 2% (w/v) uranyl acetate. Micrographs were taken under low dose condition on a JEOL JEM-1230 LaB6 transmission electron microscope operated at 120 kV with an Ultrascan 4000 SP CDD camera. Images were recorded with a nominal magnification of 10,000 (1,1 nm per pixel) per sample.

### 2.5. Biological Assays of IL-18 and IFNa-2a Variants

*Proliferation assay.* Daudi cells (ATCC CCL-213) were cultured in RPMI medium supplemented with 25 mM HEPES, 1 mM L-glutamine, 10% fetal calf serum (FCS) and 1% penicillin/streptomycin (pen/strep). Daudi cells were labelled with carboxyfluorescein succinimidyl ester (CFSE, C34554, ThermoFisher Scientific) in phosphate-buffered saline

(PBS) at a concentration of 2 nmol/40*$10^6$ cells in accordance with manufacturer's instruction. Subsequently, 200.000 cells/well were seeded in 24 well plates and they were allowed to rest for 3 h at 37 °C and 5% $CO_2$. Half-logged concentrations of commercial, heterologously expressed and glycan-tagged IFNα-2a variants (1000 to 0.01 pM) were added to the cells and compared to commercially purchased IFNα-2a (C11100, PBL Assay Science). Cells were harvested for flow cytometry analysis after 5 days and dead cells were excluded using a live/dead marker for 405 nm excitation (L34955, ThermoFisher Scientific). Acquisition took place on a CytoFlex LX (Beckman Coulter GmbH) and data analysis was performed with FlowJo, version 10.9.0 (BD Biosciences).

*IFN-γ release assay.* Human peripheral blood mononuclear cells (PBMCs) were isolated using a Ficol gradient. PBMCs were resuspended to a concentration of 2.4*$10^6$ cells in RPMI 1640 medium supplemented with 10% heat-inactivated FCS, 25 mM HEPES, 1 mM L-glutamine, 1x non-essential amino acids, 1 mM sodium pyruvate and 1% pen/strep. A volume of 250 µL were transferred into 48 well plates (0.6*10^6/well) and stimulated with glycan-tagged IL-18 variants ranging from 0.5 to 50 nM in presence of 10 ng/mL IL-12 (130-096-704, Miltenyi Biotec). PBMCs were allowed to secrete interferon-γ (IFN-γ) for 16 h at 37 °C and 5% $CO_2$. IFN-γ release was quantified by enzyme-linked immunosorbent assay (ELISA, DY285B, R&D Systems) in accordance with manufacturer's instructions. For inhibition experiments, 10 nM of each IL-18 variant was supplemented with IL-18 BP at a concentration of 0.5 and 10 nM.

*Statistical analysis.* Statistical analysis was performed with GraphPad Prism, version 10.0.2 (GraphPad Software Inc.). Statistically significant differences between groups were evaluated with the non-parametric One-way ANOVA test (Friedman test). Curve fitting was performed and the $EC_{50}$/$IC_{50}$ values were evaluated using the sigmoidal dose response function with constrained hill slope.

### 2.6. Biological Assays of Insulin Variants

*Cell culture and treatment.* SGBS pre-adipocytes were cultured at 37 ºC in Dulbecco's Modified Eagle Medium (DMEM)/F12 containing 10% FCS. Confluent cells were induced to differentiation using the following scheme (Figure S15A). Differentiation was started (day 0) by washing cells 3 times with PBS and then changing to a serum- and albumin-free differentiation medium (DMEM/F12 supplemented with 2 µmol/l rosiglitazone, 25 nmol/l dexamethasone and 0.5 mmol/l methyl3-isobutyl-1-methylxanthine). After 4 days, medium was changed, and cells were further cultured in DMEM/F12 supplemented with 0.1 µmol/l cortisol, 0.01 mg/mL transferrin, 0.2 nmol/l triiodothyronine, and 20 nmol/L human insulin. After 14 days of differentiation, cells were serum-starved for 16 h and insulin signaling was assessed following an acute stimulation with either human insulin (Sigma-Aldrich) or glycovariants **9a-9d** (5 and 100 nM or vehicle (PBS)) for 5 min. Cells were then lysed in a buffer containing 10% (w/v) glycerol, 3% (w/v) SDS and 100 mM Tris–HCl (pH 6.8). Lysates were immediately boiled for 5 min and centrifugated (13.200 rpm for 2 min). Protein content of the supernatant was determined using a bicinchoninic acid protein assay kit (23225, ThermoFisher Scientifics) (Figure S15B).

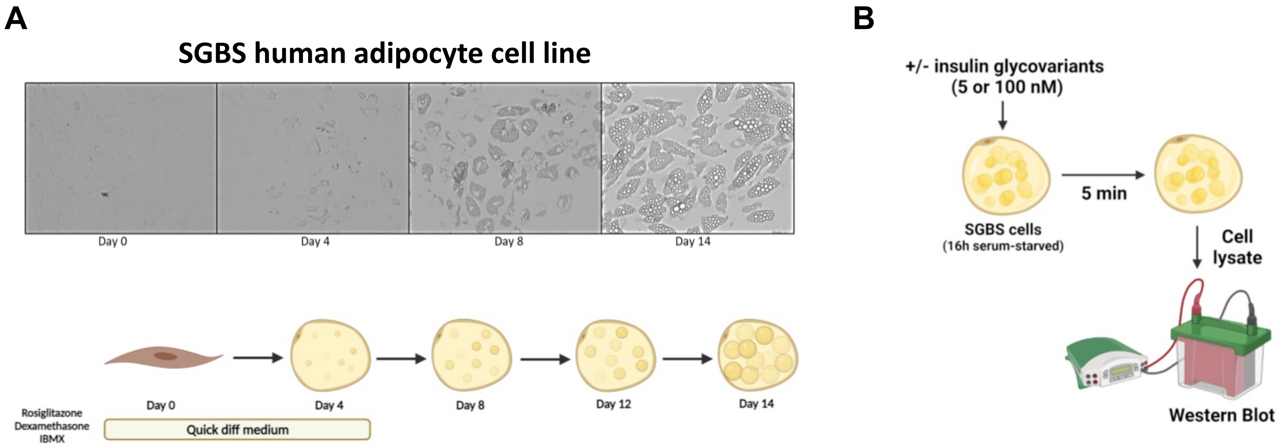


**Figure S15.** Experimental set-up for downstream signalling analysis upon administration of human insulin 9 and glycoinsulin variants 9a-d.

*Western blot analysis.* Proteins (10 µg) were separated by 10% SDS-PAGE followed by transfer to a PVDF transfer membrane. Membranes were blocked for 1 h at RT in tris-buffered saline buffer containing 0.05% Tween-20 with 5% non-fat dry milk followed by an overnight incubation with specific antibodies. Blots were then incubated with horseradish peroxidase-conjugated secondary antibodies for 2 h at RT. Antibodies against PKB/Akt-Thr308 (#9275), PKB/Akt-Ser473 (#4051), and phospho-PKB substrate (#9614) were purchased from Cell Signalling Technology (Danvers, MA, USA). Antibodies against total PRAS40 (AHO1031) and phospho-specific PRAS40-Thr246 (44-1100G) were purchased from Invitrogen (Carlsbad, CA, USA). Antibody against b-actin (A5441) was purchased from Sigma-Aldrich.

*Statistical analysis.* Graphs and statistical analyses were performed using GraphPad Prism, version 10.0.2 (GraphPad Software Inc.). All data are expressed as means ± standard error of the mean (SEM). Statistical significance was evaluated using was tested with a mixed effect model using Restricted Maximum Likelihood (REML) for data fitting (implemented in Prism 10).

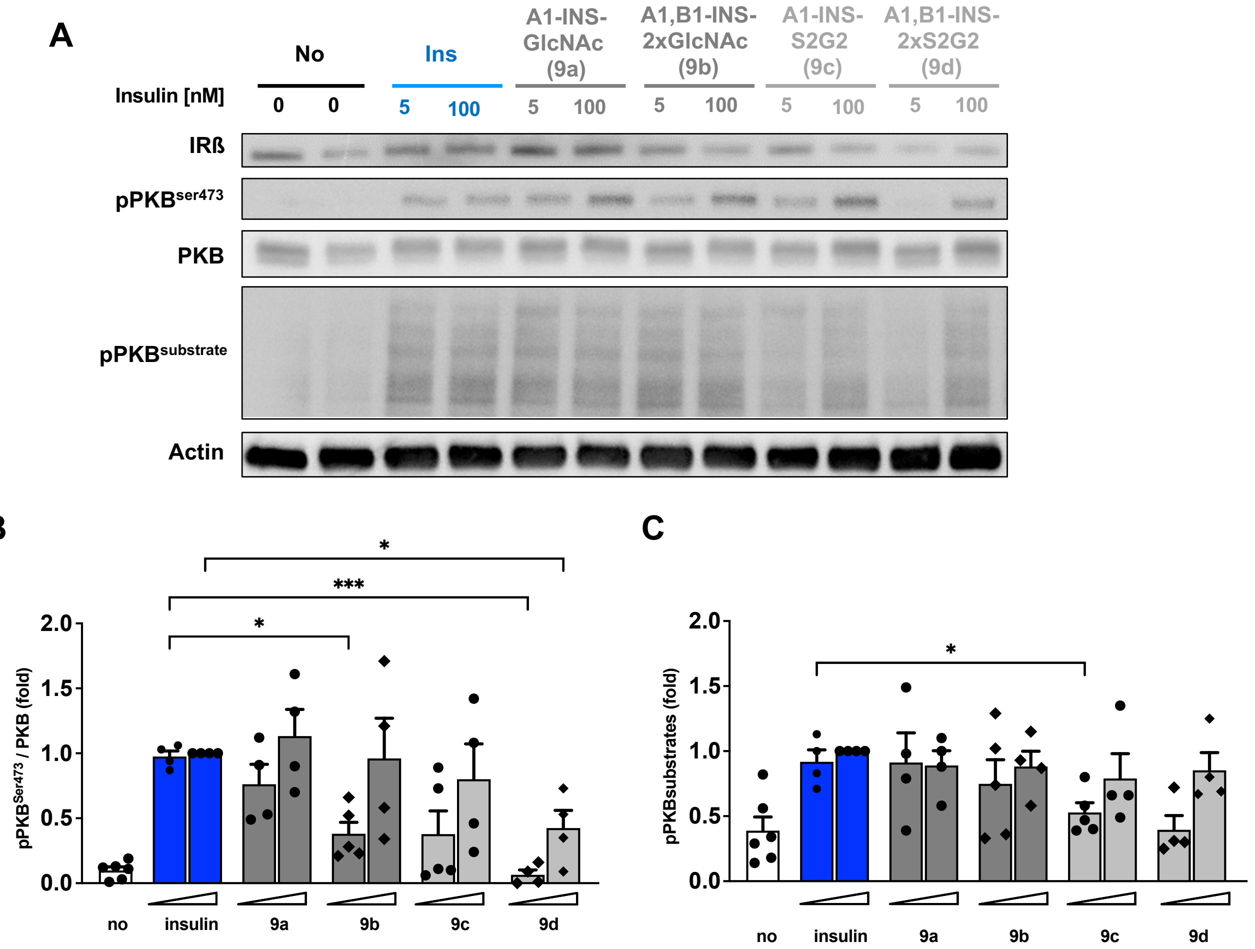


**Figure S16.** Insulin signaling by human insulin 9 and glycoinsulin variants **9a-d**. SGBS human adipocyte cells (n=4) were differentiated for 14 days in adipogenic medium and before stimulation with human insulin and glycovariants **9a-d**, the cells were serum-starved for 16 hours. Cells were lysed upon stimulation for 5 min with 5 and 100 nM of the respective variants. A) Phosphorylation of PKB (Ser473) and PKB substrates was analyzed by Western Blot. B) Phosphorylation levels were quantified for the phosphorylation site Ser473 of PKB and normalized against the non-phosphorylated protein. C) Quantification of phosphorylation levels of pKB substrates. Data are represented as means +/- standard error of the mean. Statistical significance was tested with the mixed effect model, which uses Restricted Maximum Likelihood (REML) for data fitting (implemented in Prism 10).
